# Constraint-based analysis for causal discovery in population-based biobanks

**DOI:** 10.1101/566133

**Authors:** David Amar, Euan Ashley, Manuel A. Rivas

## Abstract

Availability of large genetic databases has led to the development of powerful causal inference methods that use genetic variables as instruments to estimate causal effects. Such methods typically make many assumptions about the underlying causal graphical model, are limited in the patterns they search for in the data, and there is no guide for systematic analysis of a large database. Here, we present cGAUGE, a new pipeline for causal Graphical Analysis Using GEnetics that utilizes large changes in the significance of local conditional independencies between the genetic instruments and the phenotypes. We detect cases where causal inference can be performed with minimal risk of horizontal pleiotropy. Moreover, we search for new graphical patterns to reveal novel information about the underlying causal diagram that is not covered by extant methods, including new direct links, colliders, and evidence for confounding. We present theoretical justification, simulations, and apply our pipeline to 70 complex phenotypes from 337,198 subjects from the UK Biobank. Our results cover 102 detected causal relationships, of which some are new and many are expected. For example, we detect a direct causal link from high cholesterol to angina and a feedback loop between angina and myocardial infarction. We also corroborate a recent observational link between asthma and Crohn’s disease. Finally, we detect important features of the causal network structure including several causal hubs such as intelligence and waist circumference.

## Introduction

Modern population-based biobanks are large datasets with extensive phenotypic and genotypic data of the same subjects. These datasets offer new opportunities for causal inference due to their size and depth. However, the genetic variants typically have weak associations with the phenotypes, which can limit statistical analysis. Methods such as Mendelian Randomization (MR) and genetic-correlation (GC), utilize these genetic variables as instruments to estimate the causal effects between traits ^1–6^. Such methods deal with the genetic instruments using a *2D analysis* of their summary statistics: given a pair of phenotypes X,Y the effects of the variants of X with both (hence, 2D) are analyzed and a summary is reported. To enable causal inference, these methods typically make strong assumptions about the underlying causal diagram (*causal assumptions*) and require a specific parametric model (*statistical assumptions*). For example, standard MR assumes linear effects and reports a summary of a linear fit in the 2D plot. The basic causal assumptions are that the genetic variants are independent of confounders that affect both phenotypes, and *no horizontal pleiotropy*: instruments of X affect Y only through X ^5^. MR-PRESSO ^2^ is a recent extension of MR that directly estimates horizontal pleiotropy by correcting for variants with outlier effects. MR methods have another limitation in that they may assume nonexistence of a directed cycle between an exposure and an outcome. A directed cycle, also denoted as a feedback loop ^7^, or reverse causation ^3^, occurs in the causal diagram when two phenotypes are causes of each other, possibly indirectly. LCV ^1^, which is an example of a GC method, assumes that there is a latent variable that mediates the genetic correlation and then compares the effects of the two variant sets to assess if one phenotype is fully or partially genetically causal for the other. LCV estimates the consistency of the variant effects and the connection with causality is based on the added assumptions to the graphical structure.

2D methods are limited to analyzing summary statistics and are thus oblivious to conditional independencies (CI) detected while analyzing individual level data. In addition, they do not offer a guide for a systematic analysis of a large database, including assessing limitations of the biobank, which phenotypes to analyze, and which of the variants to take as instruments. The causal graphical models literature offers a more general framework for causal discovery that utilizes CI tests even when less causal assumptions are made ^8,9^. Such analyses are fundamentally different when compared to standard statistical analysis in that they are based on finding patterns in the data that exclude possible causal models instead of having a single objective function to be optimized (e.g., maximum likelihood). As a result, such analyses may provide partial information about causal relationships, but the general framework is more reliable when analyzing observational data ^8^.

Here, we leverage this theoretical framework to propose conditional independence (CI) methodology for causal discovery in a population-based biobank ^7–12^. We detect cases that are likely to fit standard MR assumptions, and use 2D methods to evaluate causal effects. We also utilize cases in which the CI test of a variant-phenotype is substantially different than the marginal association of the variant with the phenotype. These are used to: (1) identify direct and indirect causal relationships, (2) detect feedback loops (i.e., causal cycles), and (3) detect problematic phenotype pairs for which the data are inconclusive and 2D methods are not expected to work well. We present theoretical justification of the proposed methods, simulations to support the way we handle weak instruments, and results from applying our tools to 70 complex traits and diseases to data from 337,198 subjects from the UK Biobank ^13,14^.

## Results

### Simulations of graphical models with weak instruments

To utilize CI tests for causal discovery, we first performed simulations to test their output when instruments are weak and causal effect sizes are low. We tested four graphical cases using standard linear models with different parameterization (**Supplementary Text**). All simulated cases followed linear relationships between the variables (e.g., the linear effect of a binary instrument on a continuous exposure was set to ɑ_1_). We computed the p-values of CI tests using linear or logistic regression. **Figure 1** shows the CI p-values of selected examples, see **Supplementary Figures 1–6** for all cases. The results can be summarized as follows. First, except for very low effect sizes (ɑ_1_<0.2, which is ≤0.5% explained variance), a directed (causal) edge between an instrument and a phenotype remained robust to conditioning, even when conditioned on the affected phenotype (i.e., still had a significant p-value, **Figure 1A-B**). Second, when the instrument was indirectly linked to a phenotype (i.e., by a directed path with more than a single edge), significant association emerged only at medium-high effect sizes (due to mediation and added noise by variables along the path). Third, when an unobserved confounder was added, collider bias emerged only when the confounding effect was large, see **Figure 1B**. Finally, when we simulated a case without a causal effect, but with a joint confounding of genetics and the phenotypes (e.g., population structure), CI tests did not reveal an asymmetric structure in the data, see **Figure 1C**.

**Figure 1.**
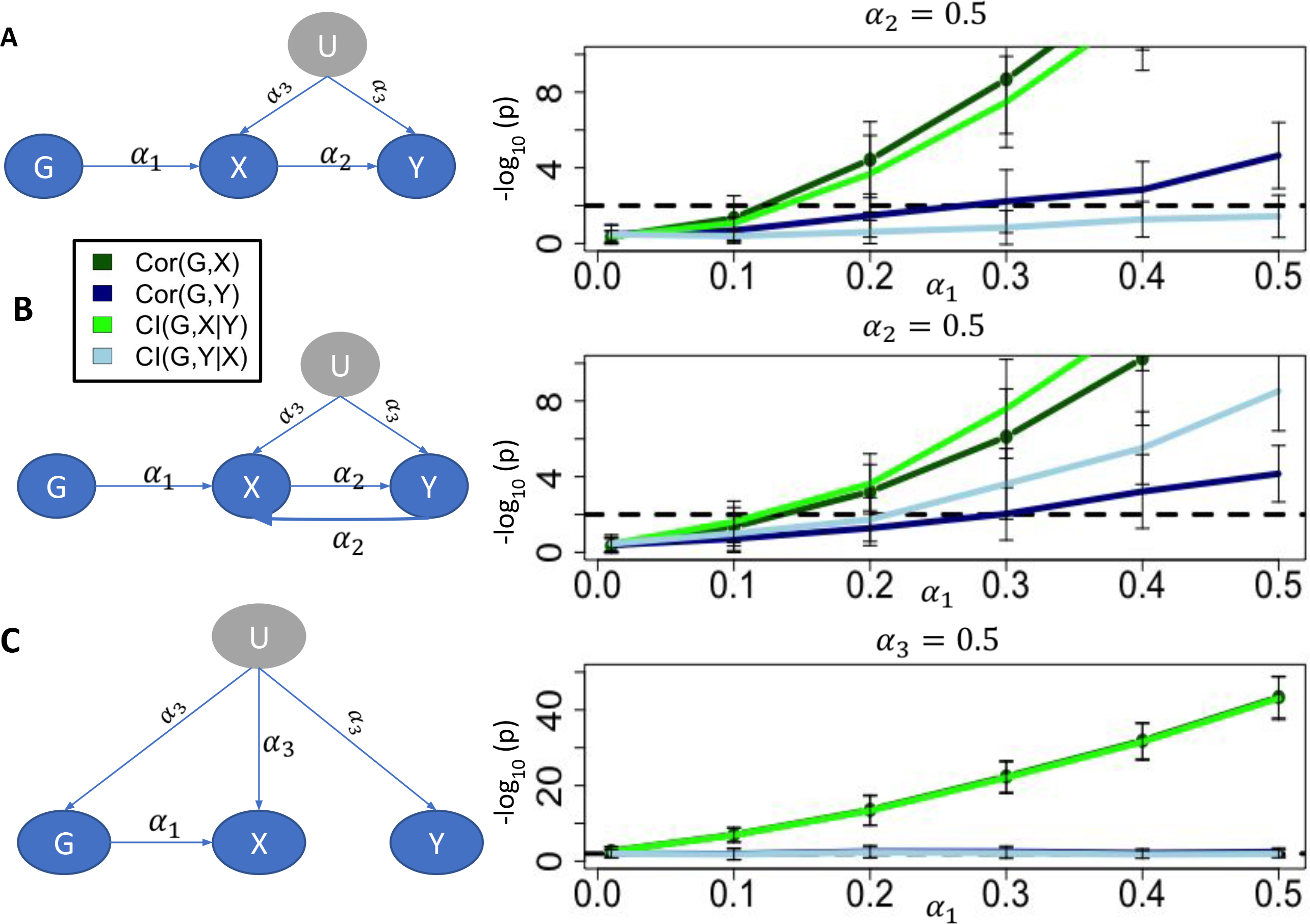
The performance of conditional independence tests in different models. G represents a binary genetic variable with 5% frequency. X and Y are two observed non-genetic continuous phenotype. U is an unobserved continuous confounder. Except for G, noise factors were set to standard normal distributions. Each bar represents the results over 100 simulated datasets. Conditional independence was tested using linear regression or logistic regression. A) The standard model assumed by MR methods. The effects of U and X on Y were set to 0.5 (< 20% explained variance). Dashed lines represent a 0.01 p-value threshold. Theoretically, conditioned on X, G and Y have an active pathway because X is a collider in the path from G to Y through U. However, this path does not manifest a detectable conditional dependence in our case as much stronger effects are needed. B) Same as (A) but now X and Y form a cycle, which means that theoretically now G and Y cannot be separated even when conditioned on U and X. In practice, compared to (A) the conditional dependencies are now stronger and detectable at lower thresholds. C) U now affects all observed variables, including G (e.g., as may occur due to uncorrected population bias). However, X has no causal effect on Y. Interestingly, in practice, when conditioned on X (Y), G and Y (X) are independent.

We were further interested in the different patterns that weak instruments reveal. We define an *emerging association* as the case in which a new association is detected between the genetic variant and the phenotype only after conditioning on another phenotype (i.e., the p-value decreases substantially, from >0.1 to < 0.001). We define a *disappearing association* as the case in which the variant and the phenotype become independent because of the conditioning (i.e., the p-value increases substantially from <0.001 to >0.1). We define a *consistent association* when the variant and the phenotype remained associated even when conditioned (p<0.001 in all tests).

Our simulations show that when the two simulated phenotypes had both a causal path and were confounded, with and without a cycle, the instruments were either not significant at any test (p>0.001) or had one of the three patterns above (i.e., emerging, disappearing, or consistent). The different patterns emerged in some cases even when the nodes were connected in the graph, see **Figure 2** and **Supplementary Figure 6**. The appearance of these patterns highlights how weak instruments of the same phenotype can have association patterns that manifest different paths even though the underlying causal diagram is fixed.

**Figure 2.**
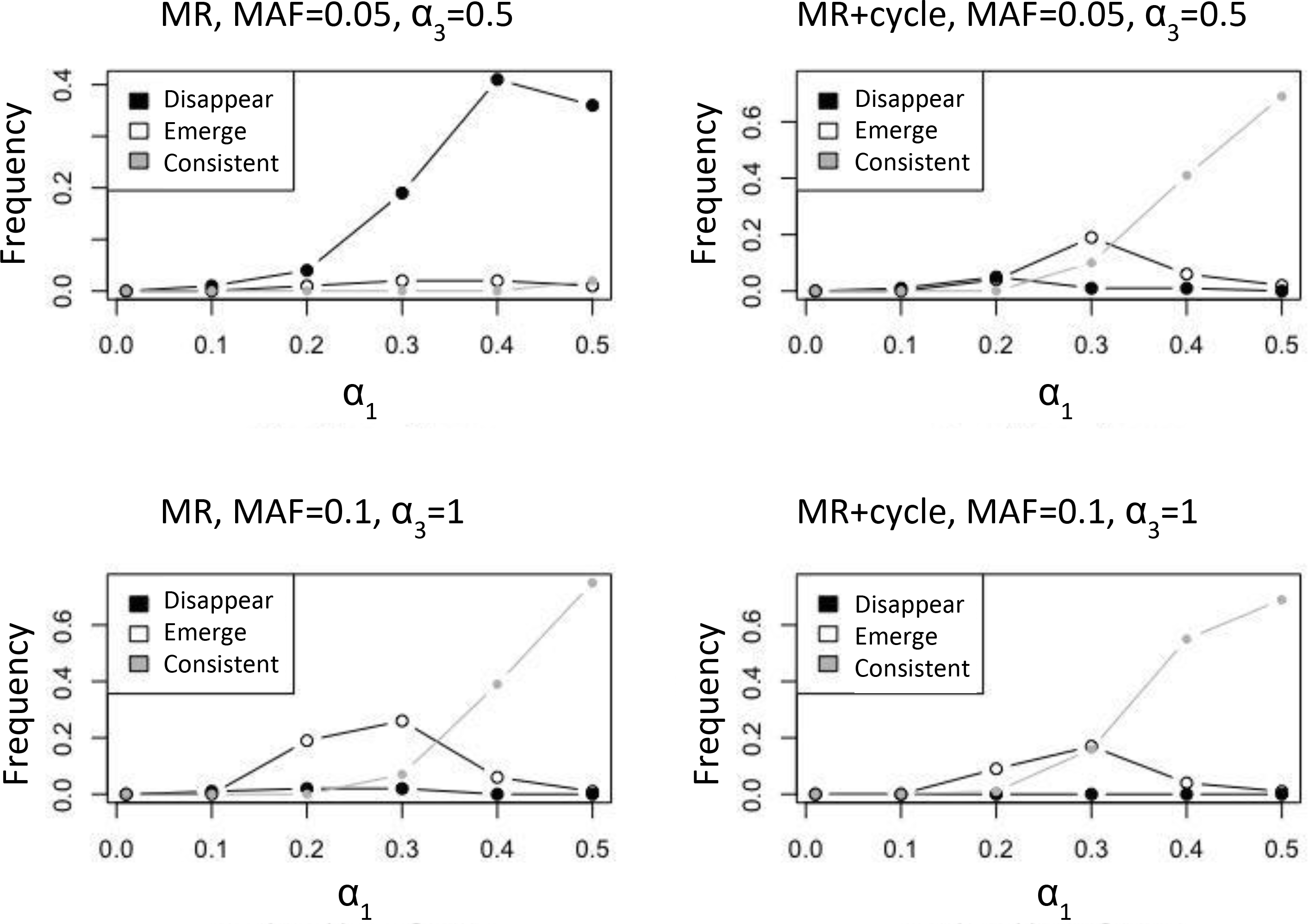
Weak instruments can provide different evidence for the same graphical models. The presented cases fit the first two simulated cases of Figure 1. Each line presents the frequency of different dependency patterns between the simulated genetic instrument G and the phenotypes Y and X. In all simulated cases G and X were significantly associated at p < 0.001, and are thus not shown. Disappearing associations: G and Y are associated marginally, but become independence when conditioned on X (p > 0.1). Emerging associations: G and Y are independent but become associated when conditioned on X. Consistent: G and Y are significantly associated with and without conditioning on X. The results show that when instruments are weak, even though the causal diagram is fixed, some may provide different patterns. Nevertheless, as expected from the theory, associations occur only when a non-blocked path exists between G and the Y.

### A novel pipeline for causal discovery using genetic instruments

Given the results above, we developed a new pipeline, cGAUGE: Causal Graphical Analysis Using GEnetics, for causal discovery using genetic instruments. See **Figure 3** for an overview, and the **Supplementary Text** for a thorough explanation, including theoretical justification. We take as input the genetic and phenotypic individual level data of a population-based biobank, a CI test (e.g., using linear or logistic regression), and two p-value thresholds: one for rejecting the null of conditional independence p_1_ and one for accepting it p_2_ (values in between are considered unreliable). Note that standard statistical tests are not designed to accept null hypotheses. However, this is a standard assumption made by causal discovery algorithms as they rely on detection of independencies ^8,9^. The phenotypic data includes exogenous variables that are adjusted for, including ethnicity, sex, age, and genetic principal components (top 5 in our case). In addition, we have a set of additional phenotypes where we perform causal discovery.

**Figure 3.**
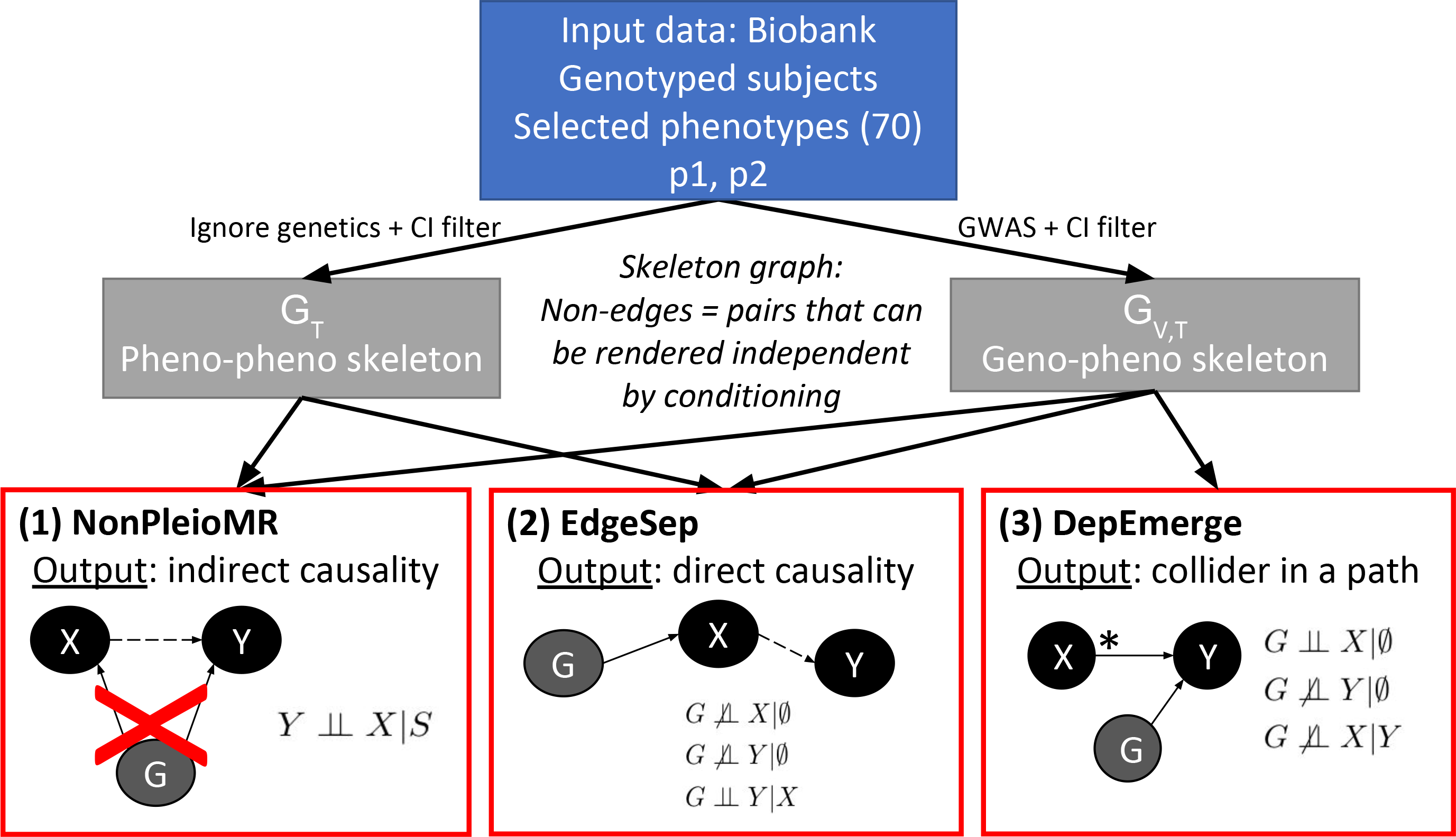
Our pipeline for causal diagram discovery using genetic data. We take as input data from independent subjects that were phenotyped and genotyped (a “biobank”). A set of phenotypes are selected in addition to standard exogenous variables such as ethnicity, sex, age, and genetic principal components. Our goal is to discover causal relationships among these phenotypes. We first preprocess the data to infer two skeleton graphs: graphs that represent associations that are robust to conditioning. Based on causal inference theory, surviving associations contain the subset of true causal links. The last layer illustrates the three graphical rules in which we utilize genetic variables to provide evidence for flow of causal information. NonPleioMR detects phenotype pairs that are unlikely to be affected by horizontal pleiotropy and run MR using their detected instruments. The instruments for this analysis are taken from the genetics-phenotypes skeleton. EdgeSep checks for associations that appear in the GWAS results of a phenotype Y (at a significance level p_1_), but disappear once conditioned on a new phenotype X (i.e., the significance is >p_2_), and such that X and Y are adjacent in the G_T_. These cases can point out a direct causal link from X to Y. Finally, DepEmerge seeks genetic variables that form new associations with a phenotype X once conditioned on Y. Such analyses hint for existence of a path in which Y is a collider and X is not.

Briefly, we first compute *skeletons* that represent associations that are robust to conditioning (i.e., significant at a predefined level in all CI tests, **Figure 3, middle layer**): one for phenotype-phenotype associations, called G_T_, and another for genetic variants-phenotypes associations, called G_I,T_. The skeletons are expected to contain the true direct causal relationships as a subset. G_T_ is composed of phenotype pairs that are significantly associated at a significance level p_1_ even when conditioned on the exogeneous variables and any pair of other phenotypes. G_I,T_ is created by first taking all standard (LD-clumped) GWAS results from each phenotype at a significance level p_1_. We then exclude (G,X) associations for which there exists a phenotype Y such that conditioning upon it (in addition to the exogenous variables) results in CI p-value>p_*2*_.

We use the skeletons and the CI tests results in three different analyses. First, *NonPleioMR* (**Figure 3**, **box 1**) is based on the observation that a phenotype pair (X,Y) that is unconnected in G_T_ cannot be under genetic confounding (horizontal pleiotropy) and thus is a suitable pair for MR. However, note that these pairs can represent indirect causal relations only. We use by default the simple MR-IVW meta-analysis to evaluate if there is evidence for X->Y ^3,15^, but any MR method can be used (e.g., MR-PRESSO). As the input for MR we take the variants that are connected only to X in G_V,T_. Using these instruments we also estimate *π*^1^: the proportion of non-null p-values (i.e., when examining the p-values of the association of the instruments with Y) under the assumption that the p-values follow a mixture distribution. Thus, this score directly measures consistency and avoids specific parametric assumptions made by 2D analysis methods (e.g., having a linear causal effect). We select as output associations that were either significant in the MR at 0.01 Bonferroni correction (of all pairs) and had >10 instruments, or pairs that had extremely high estimated *π*^1^ (>0.8).

Second, *EdgeSep* (**Figure 3**, **box 2**) takes an edge (X,Y) in G_T_ and seeks associations that appear in the GWAS results of a phenotype Y at a significance level p_1_, but disappear once conditioned on a new phenotype X (p>p_2_). For each phenotype pair we report the number and percentage of GWAS-associated genetic variables that satisfy the condition above. These patterns point out direct causal relationships. This analysis can find evidence for both X->Y and Y->X, which indicates the existence of a cycle (i.e., a feedback loop).

Finally, *DepEmerge* (**Figure 3**, **box 3**) seeks genetic variables that form new associations with a phenotype Y once conditioned on X. This serves as partial evidence for the existence of a path in which X is a collider and thus not a cause of Y. For each X,Y we report the number of variants and the percentage of the instruments of Y that fit the DepEmerge pattern. When DepEmerge discovers many variants for both (X,Y) and (Y,X), it suggests the existence of either a latent confounder for both or a cycle between them. When this occurs, these two options are *unidentifiable*: there is no statistical method that can distinguish the two cases unless additional assumptions are made ^8^. In addition, the fact that many of the instruments of one are not marginally associated with the other, it suggests that MR will not work well in practice. Thus, while the output of DepEmerge is both partial and harder to interpret, it is very valuable in that it reveals pairs of phenotypes on which 2D methods may be unreliable or fail to detect expected causal links (based on the theory). This cannot be solved, unless the researcher either has additional information or is willing to make additional assumptions (e.g., by selecting variants using external information or by handling reverse causation of diseases by excluding their subjects, see ^3^).

### Robustness analysis and comparison with other methods

We compared cGAUGE with state of the art methods: MR-Egger ^16^, MR-IVW, MR-PRESSO, and LCV. It is important to note that cGAUGE is not directly comparable with these methods both in that it is not based on summary statistics alone (i.e., 2D analysis) and that its focus is on discovery of existence of causal relationships and not inference of effects. Nevertheless, we compared the methods in the number of inferred links and their robustness. We used 70 phenotypes from 337,198 genotyped subjects, see **Supplementary Table 1**. The robustness analysis was performed by randomly splitting the dataset into two parts of the same size, running the pipelines of each subset, and comparing the detected networks using the Jaccard coefficient of their edge set, see **Materials and Methods**.

When testing different parameters for cGAUGE (p_1_=1e-8,1e-7,or 1e-6; p_2_>0.1,0.01, or 0.001, **Supplementary Table 2**), we discovered an overall high robustness of EdgeSep and DepEmerge (median J>0.72, even though they may rely on a small number of genetic instruments), and a lower robustness for NonPleioMR (J<0.45). Comparing cGAUGE to other methods (**Supplementary Table 3**) revealed that they were less reliable markedly in one of two ways: (1) when the number of output edges was moderate (<160, MR-Egger) or low (<20, LCV) the Jaccard score was low (J<0.46 for MR-Egger; J=0.33 for LCV), or (2) when the number of edges was very high (>370, MR-IVW or MR-PRESSO) the Jaccard score was high but the output was likely inflated with false positives such as multiple cancer-cancer inferred causal relationships (e.g., skin->bladder). In summary, cGAUGE offered a better balance between the number of detected associations (90-130), robustness, and reliability.

### Results on the UK Biobank data

We take as default p_1_ = 1e-07 and p_2_ = 0.001 as it gave a reasonable tradeoff between number of edges and robustness in our analysis above. Moreover, it fits both best practices from MR publications for p_1_^17,18^, and default settings of causal discovery algorithms for p_2_^19^. Nevertheless, using other combinations gave similar results and had similar scores in the robustness analysis. The results using p_1_ = 1e-06 and p_2_ = 0.001 are shown in **Supplementary Figures 7–9**.

*Skeletons*. **Figure 4** shows the inferred G_T_. This graph is sparse with 249 edges for 68 nodes, yet related phenotypes form dense regions. For example, even though the graph is one large connected component, heart diseases and their related phenotypes formed a dense module that had no behavioral phenotypes. For inference of G_V,T_ we observed that 42.5% of the original GWAS results were filtered, illustrating how GWAS results can be substantially filtered when many phenotypes are analyzed together using CI tests. From the point of interpreting a GWAS, this filter’s output can be thought of a way for prioritizing the discovered loci.

**Figure 4.**
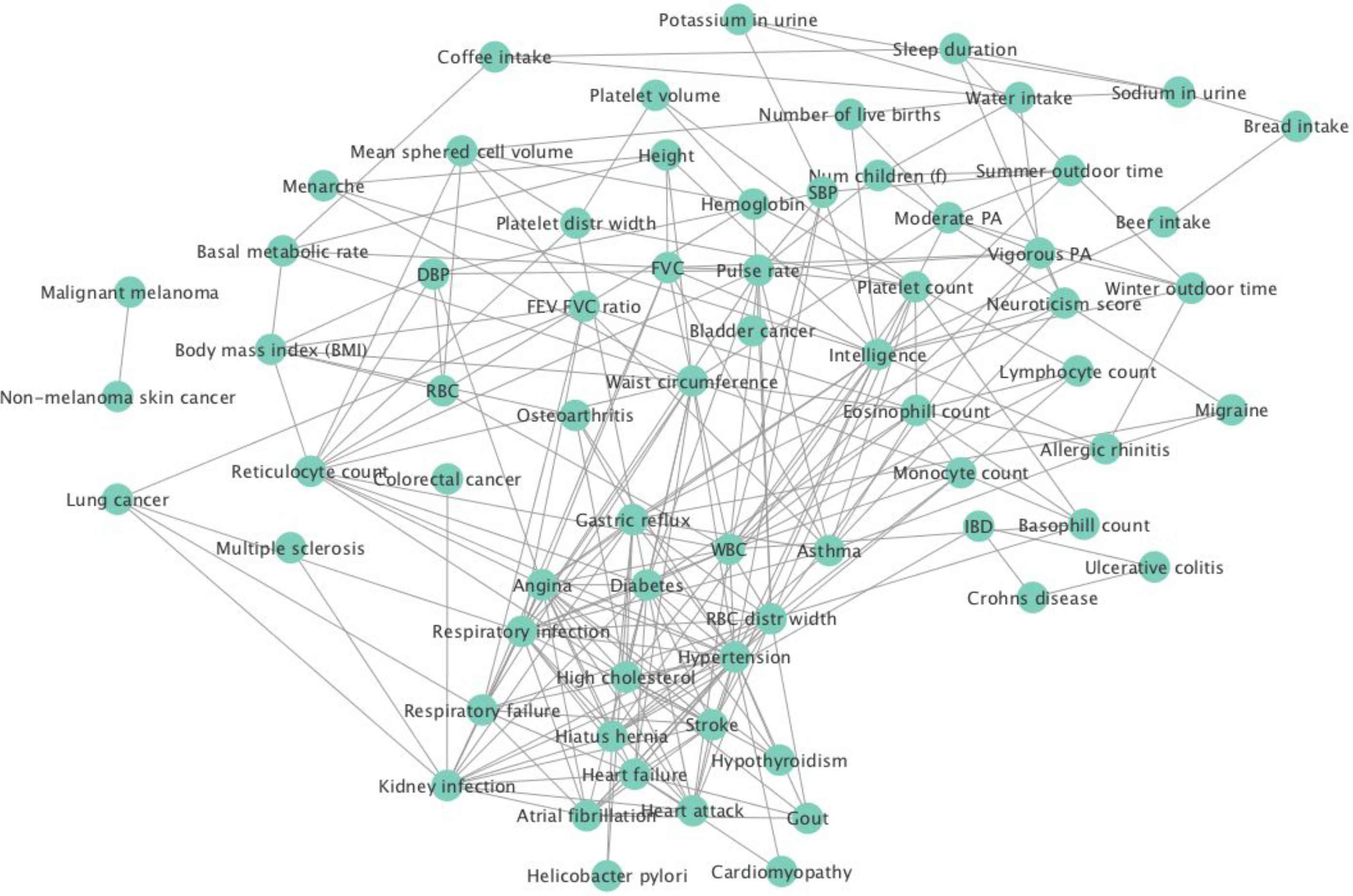
Inferred skeleton among the phenotypes (G_T_). The edges represent phenotype pairs that remain associated at p<1e-7 when conditioned on other phenotypes. The graph structure points out expected modules such as the dense area around heart attack.

*NonPleioMR* detected 24 causal links of which most are expected and some are novel, see **Supplementary Table 4.** For example, we detect asthma->Crohn’s disease, which has been hypothesized recently in an independent study ^20^. Interestingly, this link had marginal significance using LCV (p=0.09), but was significant when we tested MR-PRESSO instead of IVW as a comparison (p<1e-20).

*EdgeSep’s* results are shown in **Figure 5** and **Supplementary Table 5**. It revealed 66 causal relationships supported by at least two variants, and 12 with a only a single variant. For most pairs the percentages are low (<2%), which may be a property of partial signal detected by weak instruments (see **Supplementary Text** for discussion). Such rare local patterns are very informative, but will be overlooked in the summary analysis of 2D methods (see the example below).

**Figure 5.**
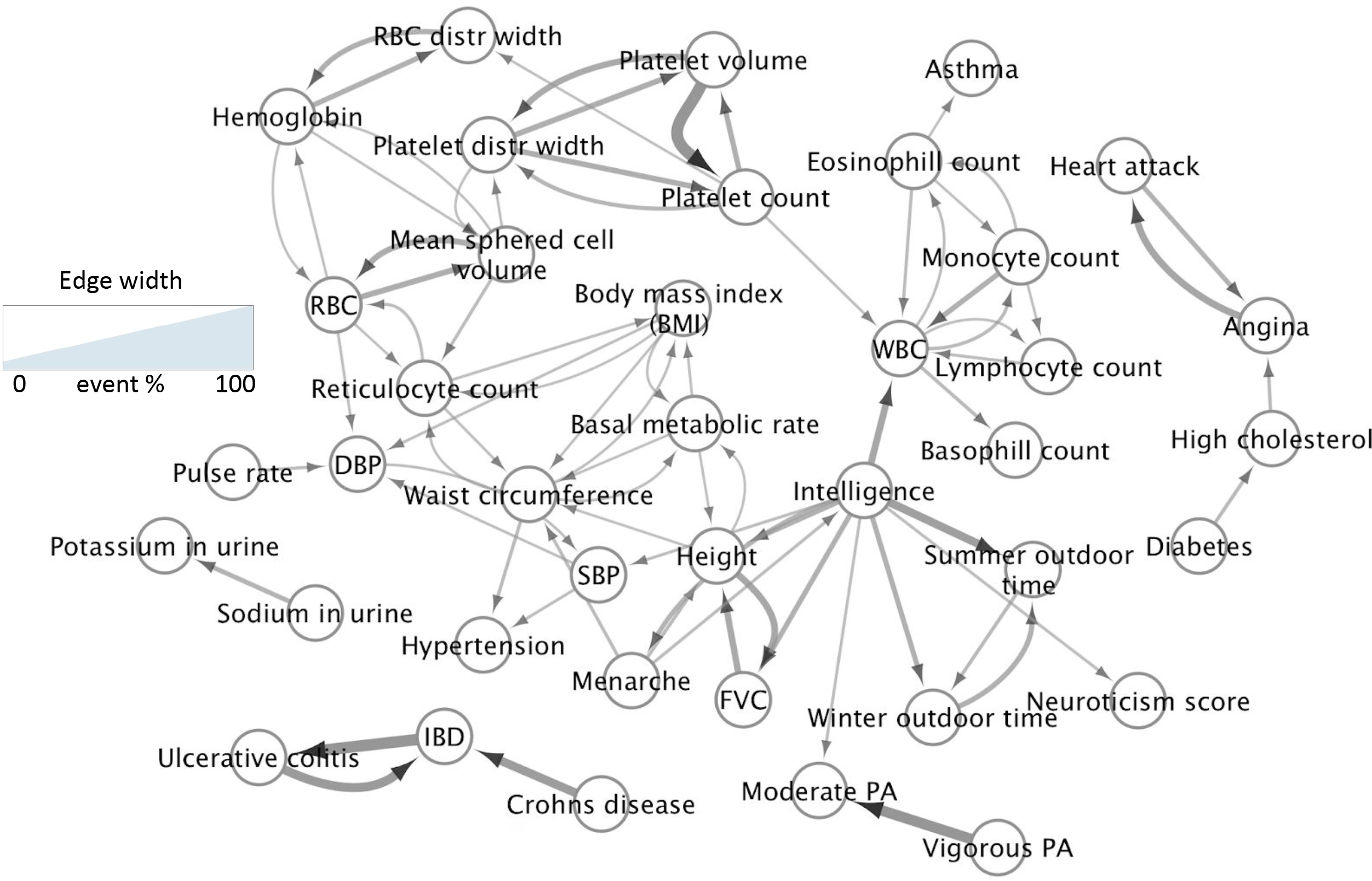
Inferred direct causal relations by disappearing associations along skeleton edges. The analysis reveals cycles, hubs, and clean directed paths. Intelligence, waist circumference, and whole blood count (WBC) are the main hubs. In this analysis, p_1_ = 1e-7, and p_2_ = 0.001.

The network has some expected causal links, such as eosinophil count->asthma, or a cycle between metabolic rate and waist circumference. Another example is the path diabetes->high cholesterol->angina<->heart attack, with the last link forming a feedback loop. Diabetes is known to affect cholesterol metabolism regardless of obesity, which fits our network ^21^. The angina-heart attack feedback is expected and supported by observational longitudinal data ^22^. Interestingly, the eosinophil->asthma link was detected using a single variant rs11743851 (out of >600 variants in G_V,T_), and was not significant using LCV or MR-PRESSO (p>0.5) even though the estimated genetic correlation was 26% (using LCV). The reverse link was detected by MRPRESSO (p=0.005), and not by LCV (p>0.5).

Note two additional features of the inferred network. First, it reveals hubs that impact many traits and diseases, most notable are waist circumference and intelligence. Second, some discovered causal relationships may result from temporal information, as was also observed by O’Connor and Price ^1^, which discussed a causal link from age of menarche to height. In our case we observed both age of menarche->height and intelligence->height, where the latter was supported by 12 instruments, (6.4% of the intelligence instruments used in this analysis). Positive correlation between height and intelligence is a well recognized phenomenon in children ^23^. This newly discovered causal link was not significant when tested using LCV (p>0.2), and had a marginal significance using MR-PRESSO (p=0.03, uncorrected).

*DepEmerge*’s results are shown in **Figure 6**, see **Supplementary Table 4** for additional results. As an example for a pattern, consider SNP rs769449, which is associated with high cholesterol (p<1e-100) but not with hypertension (p > 0.9). However, once conditioned on high cholesterol a new association with hypertension emerges (p=1.43e-8), suggesting the existence of a path from rs769449 to hypertension in which high cholesterol is a collider. Moreover, based on G_I,T_, rs769449 seems to directly additional phenotypes including erythrocyte distribution width.

**Figure 6.**
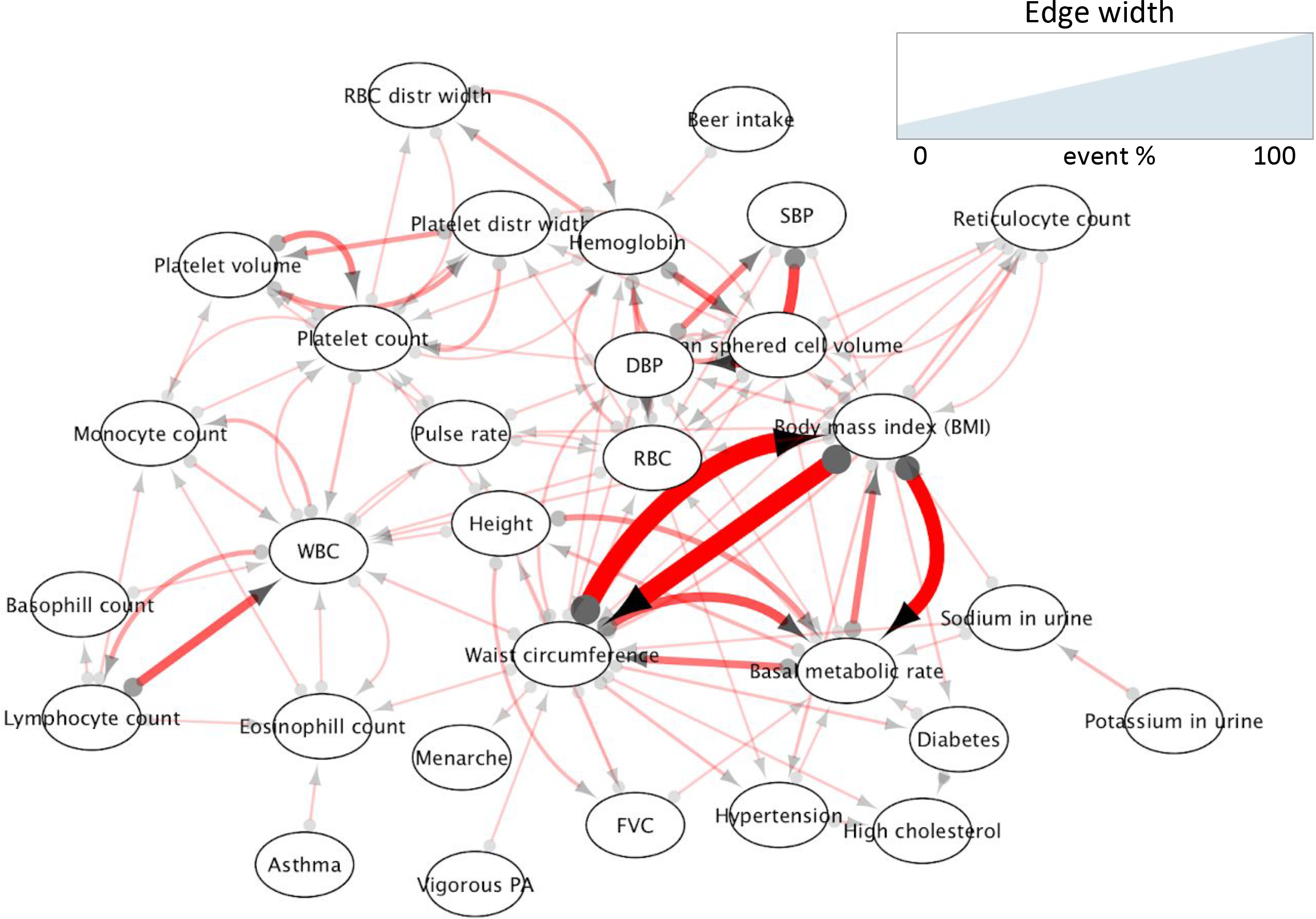
Inferred partial causal information with p_1_ = 1e-7, and p_2_ = 0.001. Each edge from X to Y represents evidence for a path without a causal effect of Y on X. When there is an edge from X to Y and an edge from Y to X it serves as evidence for latent confounding or a feedback loop. Most edges represent weak signal as the edge consistency scores are typically lower than 1%. However, in some cases (e.g., BMI vs. waist circumference) the scores are very high (> 50%). For such bi-directed edges with high percentages DepEmerge explains why MR may fail for phenotype pairs that are expected to have causal relationships.

DepEmerge’s network had most edges with low percentages but some with high (>25%), which were often bi-linked. For example, waist circumference and BMI were bi-linked with >50% support of the edges. As discussed above, we do not expect 2D analyses to work well in these cases. Nevertheless, the presence of these phenotypes as hubs in both DepEmerge’s and EdgeSep’s output suggests that these are general proxy scores for health and are thus implicated in multiple cycles. As another example for phenotypes that are present in both networks, consider asthma->eosinophil count. Our analyses suggest both a directed causal link (**Figure 5**) and a confounding effect by a third variable that affects asthma and eosinophil count (**Figure 6**), possibly medication- or behavior-related.

## Discussion

In this work we presented methods that utilize CI tests to enrich the causal discovery tool set of genetic biobanks. We filter out a substantial fraction of the single phenotype GWAS results, highlighting which genetic instruments to use. We reveal phenotype pairs with minimal violations of assumptions of MR analyses (NonPleioMR). We also search for alternating associations between genetic instruments and phenotypes to provide additional information not covered by extant approaches. These analyses both revealed features of the causal network and highlighted limitations of the analyzed dataset (EdgeSep, DepEmerge). Discovering such CI patterns is a unique property of our flow that is not covered by extant approaches such as LCV or MR-PRESSO. Other properties that distinguish our flow from such 2D analyses are: (1) we integrate the genetic information with patterns of the phenotypes inferred while ignoring genetic data, (2) we utilize patterns with provable properties under less assumptions (e.g., correctness), and (3) we detect limitations of the biobank including phenotype pairs on which MR may be biased.

In light of the points above, we view our pipeline as a preliminary analysis on which MR and GC methods can be used with better justification for either the instruments used or selection of phenotype pairs. Nevertheless, we have a few limitations that can be addressed by future studies. First, we require individual level data which may be difficult to retrieve in some cases or result in slow running times. Second, in this study we sought a proof of concept for the usefulness of graphical analysis for genetic data and reported the network of each procedure separately. Further research can build upon our flow to propose novel methods that integrate the different inferred networks. Third, we do not directly quantify the false discovery rate of an inferred network. Finally, the output of DepEmerge provides only partial information that requires additional, possibly manual interpretation.

## Materials and methods

### UK Biobank data

We used 805,462 directly genotyped variants from 337,198 white British subjects from the UK Biobank ^13,14^. The MHC region was excluded. Data were preprocessed as explained in ^24^ with a small change: we excluded variants with a MAF < 5%. 70 phenotypes (traits and diseases) were selected for the analysis, see **Supplementary Table 1**. These were selected to cover the phenotypes analyzed by ^1^, but additional traits that had large sample sizes were added.

### Single GWAS

Genome wide association analysis per phenotype was performed using PLINK ^25^. The baseline results for each each GWAS were adjusted for sex, age, and the top five genetic PCs. We also clumped the results using PLINK’s greedy approach with default parameters.

### Graph visualization

All networks were plotted using Cytoscape ^26,27^.

### Meta-analysis and Mendelian Randomization in NonPleioMR

Our pipeline discovers cases in which pleiotropic effects are unlikely (or negligible). That is, we find pairs of phenotypes (X,Y) and a set of genetic variables for X that can be used in an MR setting to test the causal effect of X on Y. However, for inference of causal diagrams alone (causal discovery), instruments can point out potential flow of information without necessarily trying to estimate effects using a parametric model. We therefore perform two meta-analyses. The first is a standard MR based on random effects meta-analysis using the IVW method ^28^. Here, we run IVW on each pair and select those whose significance passed a Bonferroni correction at 0.01. In addition, we examined the heterogeneity of the meta-analysis using Cochran’s test and excluded cases with p<1e-4 ^29^. Although this original analysis was developed to quantify pleiotropic effects, we use it in our study as a measure of statistical heterogeneity as is done in standard meta-analysis. Moreover, we observed that MR was not robust when we split the dataset (see next section) unless we added a constraint to include only analyses with 10 variants or more.

In the second analysis we simply examined the p-value distribution of the variants with the outcome variable. We computed the proportion of non-null p-values as a measure of consistency. This measure is commonly used by FDR methods and it is estimated by comparing the observed distribution of p-values to a random uniform distribution. Specifically, we use the local FDR method implemented in limma ^30,31^. We used this statistic to add causal links between pairs for which this proportion was >0.8. Our goal here was to point out cases that may still be consistent in the way the variants were associated with the exposure but were not well approximated using the MR parametric analysis.

### Other 2D analysis methods

We tested current state of the art methods by taking the GWAS summary statistics of each phenotype (without the MHC region) and ignoring the CI tests. For MR, given a p-value threshold (p=1e-6,1e-7, or 1e-8) we select the top GWAS regions with MAF >5%, clump them using PLINK, and use the filtered results as the instruments. We used the MendelianRandomization R package ^28^ to run MR-IVW, and MR-Egger. We used the MR-PRESSO implementation from the original publication ^2^. We tested MRPRESSO with and without outlier correction, denoted as MRPRESSO1 and MRPRESSO2, respectively (in Supplementary Table 3). For genetic correlation analysis using LCV we take the summary statistics (t-statistic or z-statistic) of variants with MAF>5% and run LCV with LDScore weights ^1^.

### Robustness analysis

To measure the robustness of our pipeline we split the UKBB into two random subsets of the same size. We then ran our entire pipeline and the other methods on each one and compared the inferred networks from our different analyses. For each network type we computed the Jaccard coefficient (size of intersection divided by size of union) between the edge sets. For our method we tested different values for p_1_ (1e-6,1e-7,1e-8) and p_2_ (0.1, 0.01,0.001) and computed the Jaccard score for each network type and for each combination of p_1_ and p_2_, see **Supplementary Table 2**.

### Availability

R implementation of cGAUGE is available at https://github.com/david-dd-amar/cGAUGE/.

### Funding

M.A.R. is supported by Stanford University and a National Institute of Health center for Multi- and Trans-ethnic Mapping of Mendelian and Complex Diseases grant (5U01 HG009080). The primary and processed data used to generate the analyses presented here are available in the UK Biobank access management system (https://amsportal.ukbiobank.ac.uk/) for application 24983, “Generating effective therapeutic hypotheses from genomic and hospital linkage data”(http://www.ukbiobank.ac.uk/wp-content/uploads/2017/06/24983-Dr-Manuel-Rivas.pdf), and the results are displayed in the Global Biobank Engine (https://biobankengine.stanford.edu). This work was supported by National Human Genome Research Institute (NHGRI) of the National Institutes of Health (NIH) under awards R01HG010140. The content is solely the responsibility of the authors and does not necessarily represent the official views of the National Institutes of Health.

## 1 Notation and background

In this section we give a brief overview of the theoretical results that provide the foundation for our algorithms. For a more thorough background on theory of causal inference see [1, 2].

### Notation

We generally use the following notations unless stated otherwise. Graphs: calligraphic uppercase (e.g., 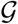). Distributions (including empirical): blackboard bold typeface (e.g., ℙ). Italic uppercase: a random variable. Italic bold uppercase: a set. Lowercase letter: a scalar or a realization of a random variable.

### Graphs

A graph 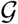 is an ordered pair 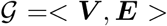, where ***V*** is a set of nodes (or vertices) and ***E*** is a set of pairs of nodes. When two nodes are connected by an edge we say that they are adjacent. In an undirected graph, ***E*** is a set of unordered pairs, whereas in a directed graph ***E*** is a set of ordered pairs and an edge < *X*, *Y* > can also be marked as *X* → *Y*. If 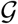 is directed and < *X*, *Y* >∈ ***E*** then we say that *X* is a *parent* of *Y*. We denote the set of parents of a node *Y* as *Pa*(*Y*). In this paper we consider directed graphs without self loops, that is *∀X* ∈ ***V***, *X* ∉ *Pa*(*X*). Generalizing parent-child relationships, we can naturally define descendants and ancestors.

An *undirected* path *π* between *X* and *Y* is a set of consecutive edges (independent of their orientation) connecting the variables such that no vertex is visited more than once. A *directed* path from *X* to *Y* is a set of consecutive directed edges from *X* to *Y* in the direction of the edges. A (directed) cycle in the graph occurs when there is a (directed) path from *X* to *Y*, and *Y* and *X* are also adjacent (i.e., *Y* → *X* in the directed case). A non-endpoint variable *X* in a path *π* is called a *collider* if and only if the edges around it have their arrowheads into *X* (i.e., *Z* → *X* ← *Y*).

### Causal models

We use the structural equation model with independent errors (SEM-IE) to describe a *causal model* [1]. This fits the standard assumptions made by current methods for genetic data analysis. Briefly, a causal model *M* is a pair 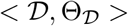 with a directed graph 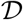 over a set of variables ***V***. 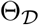 is a *parameterization* that assigns a function *x*_*i*_ = *f*_*i*_(*Pa*(*X*_*i*_), *∊*_*i*_) for each *X*_*i*_ ∈ ***V***, where *∊*_*i*_ is an error term (also called disturbance) distributed independently of other error terms according to a distribution *P* (*∊*_*i*_). Given all variables and distributions in a causal model, observed data follow a joint distribution ℙ.

### DAG factorization

The causal diagram 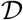 is the underlying graph that represents the direct causal interactions between the variables represented by the vertices ***V*** (some may be unobserved). If 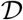 is directed and acyclic (DAG) we say that a distribution ℙ satisfies that *Markov property* if it factorizes over 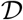 such that 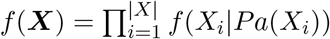 where *f* marks density functions. A similar definition can be used for discrete variables using probability functions instead of densities.

### D-separation

Early work had established that some conditional independencies can be inferred from the graph structure alone regardless of the parameterization of the entire causal model [1]. When found, such discoveries constraint the set of possible distributions that are compatible with the underlying causal structure. The graphical rule is called *d-separation* and it is defined as follows. A set of variables ***Z*** (possibly an empty set) d-separates a set of variables ***X*** from another set ***Y***, where ***X***, ***Y***, and ***Z*** are all disjoint, if and only if every path from a node in ***X*** to a node in ***Y*** is blocked by ***Z***. A path *π* between two nodes *X* and *Y* is blocked by a set of nodes ***Z***, *X*, *Y* ∉ ***Z*** if and only if at least one of the following is satisfied: (1) ***Z*** contains a node *m* that emits an edge in *π* (i.e., there is an edge *m* → *m*′ in *π*), (2) there is a collider *X*′ in *π* such that not *X*′ nor any of its descendants are in ***Z***.

We use 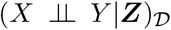 to mark the event that ***Z*** d-separates two nodes *X* and *Y* in the graph 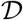. We use (*X* ⫫ *Y*|***Z***)_ℙ_ to denote the case in which two random variables *X* and *Y* are conditionally independent given a set of variables ***Z*** according to their joint distribution ℙ. The d-separation property of DAGs can be succinctly written: if 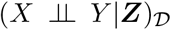 then (*X* ⫫ *Y*|***Z***)_ℙ_ in any distribution ℙ that is compatible (i.e., satisfies the Markov property) with 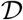.

### D-separation in graphs with cycles

Although originally proven for causal models with DAG diagrams, the d-separation criterion was later proven for diagrams with cycles in two cases: (1) discrete finite variables in positive distributions [3], and (2) linear SEM-IEs [4]. Of note, the Markov property does not necessarily hold when cycles are introduced. Therefore, additional assumptions must be used. For example, in both cases above, the proofs require the assumption that the observed distribution is an *equilibrium*. Roughly, an equilibrium in this context is reached from an SEM when we start with a random sample of the errors and then simulate the SEM repeatedly until we achieve convergence. The set of values achievable in this process is called the *equilibrium distribution*. For the linear case Fisher’s fixed point method can be used to simulate the equilibrium distribution from an SEM-IE. In the discrete case, stronger assumptions are required as the process depends on the order of the equations [5]. Nevertheless, once the equilibrium is well-defined, d-separation holds [5, 6].

### Causal discovery

The tight connection between causal diagrams and conditional independencies has been used to propose causal discovery algorithms. For DAGs, we used the *Markov condition* mentioned above, which also means that each variable *X* is independent of its non-descendants given its parents. For diagrams with cycles, while this property does not hold, the *d - separation* rule is still used. However, two additional fundamental assumptions that seem plausible in practice and can be treated as axioms are used as well [2]: *minimality*, and *faithfulness*. Minimality is the causal inference analog to Occam’s Razor: we prefer simpler models when we consider alternatives that can explain the data. More formally, a causal diagram 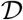 satisfies the minimality condition with respect to a distribution ℙ if the Markov property does not hold for every 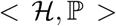, where 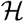 is a proper sub-graph of 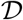. Finally, faithfulness, (also called stability) assumes that all independencies embedded in the observed distribution ℙ are stable and are invariant to changes in parameterization. Thus, it implies that 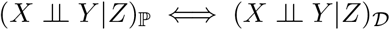.

The assumptions above are especially important if we are to propose causal discovery approaches in the present of confounding by latent variables. We generally say that the effect of *X* on *Y* is *confounded* if there exists a third variable *U* such that *X* ← *U* → *Y*. If *U* is unobserved the standard notation is *X* ↔ *Y*. We use 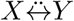 instead to avoid confusion with having a cycle *X* → *Y* and *X* ← *Y*. Presence of unobserved confounding adds complexity to the inference problem and limits our discovery power. Nevertheless, conditional independencies inferred from observed distribution are still powerful enough to constraint the set of plausible causal diagrams. When enough constraints are accumulated we can partially or completely infer causal diagrams.

It is important to note the fundamental difference between this algorithmic approach and standard statistical inference algorithms that optimize a global objective function such as training a Bayesian network using maximum a posteriori (MAP) or maximum likelihood (MLE) objectives. Here, the basic assumptions and the theoretical results are used as constraints in order to narrow down the set of possible causal diagrams. Thus, these algorithms heavily rely on the ability to statistically detect conditional independencies, and their output may be an only partially oriented diagram. Notable algorithms to perform such analyses are PC [4], IC* [1], FCI [2, 7], and CCD [8, 2]. Another notable example is the SAT-based optimization algorithm of [9] that extended the basic ideas to address the inference task as an optimization problem.

A few notable drawbacks of the algorithms above should be highlighted, especially for the case of analysis of large biobanks. First, these algorithms require exponential running times as results from all conditional independencies may be required. Second, most of these algorithms assume that the input contains a statistical oracle for testing conditional independence. Alternatively, this assumes that in practice there are no statistical errors, although some recent approaches had attempted to mitigate this issue by addressing the FDR of inferred edges [10, 11]. Nevertheless, the sequential nature by which these algorithms update the inferred structure implies that errors are propagated and accumulated. These two issues are crucial when biobanks are considered as we may need to analyze data from thousands of weak genetic variants and dozens of phenotypes. Our algorithms below build upon the main principles of these methods, integrating them with prior biological information about the nature of genetic variables.

### Instrumental variable analysis

A special set of causal inference algorithms can be defined once *instrumental variables* are present. A variable *Z* is *instrumental* relative to (*X*, *Y*) if it is not independent of *X* and is independent of all variables that have influence on *Y* that is not mediated by *X* [1]. Using instrumental variables, we can use path-analysis methods to quantify causal effects associated with the edges of an SEM [12, 13]. In the genetic and epidemiological literature such analyses are often called *Mendelian Randomization* (MR), and the recent availability of genetic biobanks has led to increased interest in new MR methods. These methods are appealing for their ability to handle many instruments, reverse causality in some situations, and confounding effects [14]. Another common approach is to compute the overall genetic correlation (GC) between phenotypes. The recently proposed LCV method [15], uses higher moments of inferred summary statistics to move beyond standard GC to causal inference under a specific parametric model.

Using genetic instruments for causal inference process is promising as they act as markers that carry causal information flow. This is a unique property not generally achievable in causal inference from observational data, which explains the high popularity of MR and GC methods. However, extant methods are limited in that: (1) they assume that the instruments are known and are not confounded themselves (i.e., by population bias), (2) they are typically based on a parametric linear model, (3) they may be sensitive to horizontal *pleiotropy*: confounding effects caused by the genetic instruments, and (4) these methods are limited to analysis of GWAS summary statistics and are therefore limited in the ability to utilize conditional independencies. In addition, there are important issues that these methods do not address such as exploring latent confounding and presenting how weak instruments behave. In our analysis we make use of constraint-based methodology to alleviate these issues and guide the causal inference process in large genetic biobanks.

## 2 Simulations: weak instruments and their effects on conditional independence tests

We evaluated the performance of conditional independence tests when instruments are weak using linear structural equations. Let *G* represent a binary genetic variable, *X* and *Y* two observed continuous variables, and *U* an unobserved continuous variable. Four different graphical structures were considered: a simple chain *G* → *X* → *Y*, and the three graphs shown in **Figure 1**. Datasets were simulated using standard normal errors for continuous variables and sampling *G* from a Bernoulli distribution with a different probability to simulate different minor allele frequencies. Then, each edge had a coefficient that determined its added value in the structural equation. For example, the graph structure presented in **Figure 1A**can be written:

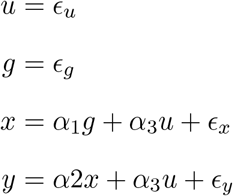

where *∊*_*g*_ ~ *Bernoulli*(*p*), and *∊*_*u*_, *∊*_*x*_, *∊*_*y*_ ~ *N*(0, 1) i.i.d.

When cycles are present, as in **Figure 1B**, we use Fisher’s iterative fixed point method [16]: we first simulate the errors and then repeatedly update the other variables, which converge almost surely to a fixed point. The convergence threshold was set to 10^−4^ or up to 100 update repeats, which were rarely required.

In all simulated cases we tested the range 0.01, 0.1, 0.2, 0.3, 0.4, 0.5 for *α*_1_, and 0.01, 0.05, 0.1 for *p*. For models without cycles we tested 0.2, 0.5, 1, 2 for *α*_2_ and omitted 1, 2 for cycles to avoid convergence issues. Finally, for *α*_3_ we considered two cases: weak confounding with *α*_3_ = 0.5, and strong confounding with *α*_3_ = 1. For each combination, we simulated 100 datasets with 10,000 samples. Conditional independence was tested using generalized likelihood tests via either linear regression or logistic regression (e.g., if *G* was considered as the dependent variable).

We were interested in the performance of conditional independence tests and their fit with the theory when weak instruments are present. We present the empirical distribution of the p-values of each test in each case. We also computed the frequencies of mixed results for each case. That is, we compared how many instruments had: (1) disappearing correlations - 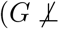, 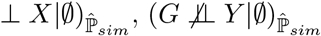, and 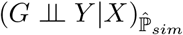, (2) emerging correlations - 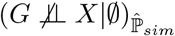, 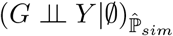, and 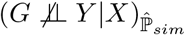, and (3) consistent correlations - 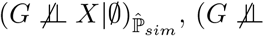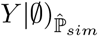, and 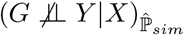. Here, 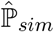, is the empirical distribution of a single simulated dataset. All simulation results are presented in **Figures 1 and 2**, and **Supplementary Figures 1-6**.

## 3 Graphical analysis to guide causal inference in genetic biobanks

### 3.1 Input data

We formalize our input as follows. We have two sets of features ***I*** (instruments), and ***T*** (phenotypes/traits/diseases) measured in *n* independent subjects. We denote the empirical distribution of the data as 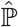. ***I*** is a set of genetic variants (typically SNPs) that are used as instruments for the causal inference process. ***T*** is a set of phenotypes, that are typically complex traits and diseases. Generally, |***I***| is large initially (>500,000), but can be substantially reduced once relevant variants are detected and LD-clumped (or pruned) per phenotype. |***T***| is much smaller. In our case we used the UK Biobank to obtain data of 337,198 white British subjects. We selected 70 phenotypes by taking those covered in [15] and some additional ones that had large sample sizes, see **Supplementary Table 1**. In addition, we assume that sex, age, and genetic principal components are exogenous variables (ethnicity as well but it was ignored here because our analysis was limited to a single group). We denote their collective set as ***W***, and assume that ***W*** ⋂ ***T*** = ∅.

Our analyses depend on three additional parameters: a test for conditional independence and the thresholds to make a decision about the results of such tests. Let *ci*(*X*, *Y*, ***Z***) be a statistical test for conditional independence (i.e., a function that returns a p-value). The *null hypothesis* of this test is that 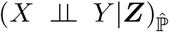, and low p-values serve as evidence for the alternative 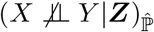. As in the simulations above, we use linear regression or logistic regression (for binary variables) for conditional independence tests. The two additional parameters are *p*_1_, and *p*_2_. When *ci*(*X*, *Y*, ***Z***) ≤ *p*_1_ we conclude that 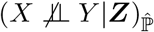. When *ci*(*X*, *Y*, ***Z***) ≤ *p*_2_ we conclude that 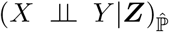. P-values in (*p*_1_, *p*_2_) represent *ambiguous* cases. Additional discussion about the importance of these parameters is given below. We tested different combinations of *p*_1_ and *p*_2_ (on real data): *p*_1_ = 10^−7^ or *p*_1_ = 10^−6^, and *p*_2_ = 0.001 or *p*_2_ = 0.01.

### 3.2 Assumptions

We first make two general assumptions about the data. First, we assume that there is no directed path going from a member of ***T*** to a member of ***I***. This assumption is biological and states that at a specific generation, the observed phenotypes do not change the DNA of the subjects. Second, we assume that the dataset is not under some collider bias. This implies that if we observe 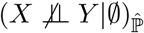 then it truly reflects 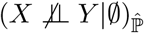 (as the sample sizes goes to infinity).

In the simulations we illustrated that even for linear models, low power can lead to *partial faithfulness fit* when analyzing weak instruments in practice: when multiple unblocked paths exist between an instrument *G* and a variable *Y*, some may manifest observed (conditional) association, and some do not. In other words, when 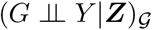 then *ci*(*G*, *Y*, ***S***) will follow the null distribution. However, if 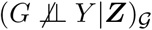 then *ci*(*G*, *Y*, ***S***) may follow some distribution that is too similar to the null distribution. Such partial faithfulness fit is very problematic for standard causal discovery algorithms as they heavily rely on a sequential process that uses faithfulness to make decisions. Thus, errors due to partial fit may propagate and distort the output.

While faithfulness may be a reasonable assumption when analyzing variables from ***T***, it is problematic when we consider weak instruments. We address this issue by introducing a relaxed assumption about conditional independence between variables from ***I*** and ***T***, which we denote as *local faithfulness of order k*, for some integer *k* ≥ 1. Standard faithfulness dictates that if *G* is a genetic instrument, *X* ∈ ***T*** (following the assumptions above), ***S***, ***S*′** ⊂ ***T*** are two sets such that *X* ∉ ***S***, *X* ∉ ***S*′**, and we observed 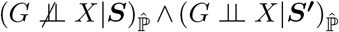 then every path in 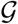 between *G* and *X* that is unblocked given *S* becomes blocked given *S*′. We alternatively assume that if |***S*** \ ***S*′**| ≤ *k* ^ |***S*′** \ ***S***| ≤ *k* and we observed 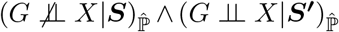 then there **exists a path***π* in 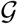 that is unblocked given *S* but is blocked given *S*′.

In words, if we observed an association between *G* and *X* given a set *S* such that a small change to *S* renders *G* and *X* independent then the paths between *G* and *X* that manifested the association before are now blocked in 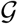 given the new set ***S***. Thus, we ease faithfulness in that we do not assume that 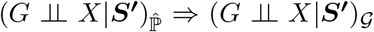 necessarily. We alternatively require some local information first that is dependent on observed patterns in the data. Note that the observed association still indicates the existence of an unblocked path between *G* and *X* given *S*, regardless of faithfulness.

Given local faithfulness we can now search for high confidence events of *alternating associations* that have provable properties that can be used for causal discovery. In this work we consider *k* = 1 only. In practice, we use information from multiple independent genetic instruments in parallel and record their patterns. In addition, as discussed above, we use two thresholds for p-values *p*_1_ and *p*_2_ that are substantially different (by orders of magnitude). That is, we seek cases in which a small change to the set conditioned upon leads to substantial difference in the association analysis result and use it as an evidence for the existence of a specific path structure in the underlying causal diagram. These analyses are explained in the next sections.

### 3.3 Algorithm overview

Our pipeline has five steps:

1. **Single GWAS**: perform a genome-wide association study for each phenotype, adjust for sex and genetic principal components (5). Clump the variants from each phenotype and merge the list.
2. **Skeletons graphs**: detect associations that are robust to conditioning.
3. **NonPleioMR**: analyze phenotype pairs that are unlikely to be under genetic confounding effects.
4. **EdgeSep**: search for skeleton edges that have evidence for a specific direction hinted by disappearing associations.
5. **DepEmerge**: search for collider bias evidence by looking at newly formed associations.

Note that unlike standard MR and GC methods we combine information from analysis of instruments with results obtained by looking at the phenotypes while ignoring the genetic data. These, together with the alternating associations detected in steps 4 and 5, are all meant to utilize the partial information that weak instruments provide for causal discovery.

For phase 1 we use standard procedures using PLINK [17]. We also avoid an exponential number of statistical tests using practical heuristics. These enables powerful and robust algorithmic strategies. The subsequent analyses are discussed below. The final section discusses meta-analysis and additional measurements that can be used to prioritize candidate causal links.

### 3.4 Skeleton graphs

Inference of a skeleton graph is the initial step that most causal discovery algorithms perform. A *skeleton graph* is an undirected graph 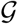 over a set of observed variables ***V*** such that an edge *e* = (*X, Y*) is added only if there is no evidence that *X* and *Y* can be rendered independent by conditioning on a set ***Z*** (which can be empty) of other observed variables. In other words, skeleton graphs represent associations that are robust to conditioning. Under our assumptions above, these graphs represent a set of candidate pairs that contain the true direct causal links in the underlying causal diagram.

As implied above, the construction of a skeleton depends on a search algorithm that scans through the space of possible sets on which we condition. Naturally, going over all possible sets is not feasible in our case. We deal with this issue using two strategies. First, we limit the number of tested sets per pair of variables to *O*(|*T*|^2^). Second, we define two skeleton graphs: 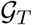 and 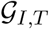. 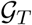 is a standard skeleton inferred by analyzing the phenotypes alone and ignoring the genetic information. 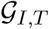 is a bipartite graph: here we examine the associations between genetic variables and phenotypes.

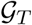 is constructed as follows. We start with a complete graph. For each pair *X*, *Y* ∈ ***T*** we go over all sets ***Z*** ⊆ *T* \ {*X*, *Y*} such that |***Z***| ≤ 2. For each such ***Z*** we compute two conditional independence tests: *ci*(*X*, *Y*, ***Z***) and *ci*(*X*, *Y*, ***Z*** ⋃ ***W***). If at least of these p-values is ≥ *p*_1_ then we remove the edge between *X* and *Y* from 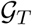.

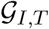 is constructed as follows. We start with an empty bipartite graph between ***I*** and ***T***. We then add an edge between a genetic variable *G* and a phenotype *X* if the GWAS result marked *G* as associated with *X* at a significant level *p*_1_. For each such *G, X* pair we then go over all sets ***Z*** ⊆ *T* \ {*X*} such that |***Z***| ≤ 1. For each such ***Z*** we compute two conditional independence tests: *ci*(*X*, *G*, ***Z***) and *ci*(*X*, *G*, **Z** ⋃ ***W***). If at least one of these p-values is ≥ *p*_2_ then we remove the edge between *X* and *G* from 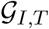.

### 3.5 NonPleioMR: Phenotype pairs that have no pleiotropic confounding

Pleiotropy is a biological phenomenon in which a genetic factor affects two or more biological processes (and their downstream phenotypes). This term is also used to describe confounding caused by genetic variables. That is, when a genetic variable causally affects two phenotypes that are investigated. We say that the pleiotropic confounding is *direct* if the effect of the genetic variable is not mediated by other phenotypes in the study (note that additional directed pathways may occur as well). Pleiotropy poses a fundamental problem for MR methods as it contradicts their basic assumptions, and therefore undermines their validity.

Another type of bias that jeopardizes using genetic variables as instruments is joint confounding with the phenotypes, as may occur due to population bias. Even though there is a rich literature that aims to deal with such biases, there is no guarantee that such bias was completely removed even when these methods are used.

We now move to using graphical guidelines to alleviate these issues.

#### Lemma 3.1.

Given the skeleton of the phenotypes 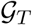 inferred using conditional independence tests among the traits only, a pair *X, Y* ∈ ***T*** that is disconnected in 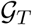 cannot have direct pleiotropic effects.

*Proof.* Had *X* and *Y* been directly affected by genetic variables then under the faithfulness assumption there could not have been a conditional independence test to separate them and thus *X* and *Y* would have been connected in 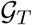, a contradiction

This lemma implies that non-edges in 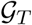 are good candidates for using MR-like methods, however it does not tell us which instruments to use, nor does it help with population bias. We address this issue next.

#### Definition 3.1.

Given the bipartite skeleton 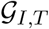 and *X* ∈ ***T***, let *Nei*_1_(*X*) be the set of all neighbors of *X* that are not linked to any other member of ***T***.

#### Theorem 3.1.

Given a pair of phenotypes *X*, *Y* ∈ ***T*** not adjacent in 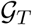, and such that |*Nei*_1_(*X*)| > 0, the variants in *Nei*_1_(*X*) are proper instruments relative to (*X*, *Y*).

*Proof.* By assumption about the correctness of 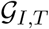, members of *Nei*_1_(*X*) cannot have outgoing edges to members of ***T*** other than *X*. Similarly, there cannot be an unobserved variable *U* (a confounder) that has pathways going to both a member of *Nei*_1_(*X*) and *Y*. Finally, by our assumption that genetic variants cannot be affected by members of ***T***, there cannot be either directed pathways from *X* or *Y* to members of *Nei*_1_(*X*), nor is there another member *Z* ∈ ***T*** that affects *X* or *Y* and is a member of *Nei*_1_(*X*)

The two lemmas above help us integrate information from both sources: instruments-phenotypes and the phenotypes’ joint distribution. When we limit our analysis to phenotype pairs that are not adjacent in 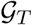 we rule out the possibility of strong unobserved confounding, pleiotropic or not, even if we have errors in 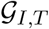. On the other hand, once these pairs has been established, the safest variant set to use as instruments are those in *Nei*_1_(*X*). Note that if we are willing to assume that 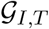 has no errors then *Nei*_1_(*X*) can be used as instruments (e.g., for MR) relative to any pair (*X*, *Y*) (*Y* ∈ ***T***, *Y* ≠ *X*). However, as a conservative measure we chose to add the constraint from lemma 3.1 as it represents compelling evidence that is independent of the genetic variables.

### 3.6 EdgeSep: Disappearing associations along skeleton edges

In the analysis above we excluded the detected edges in 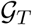. These edges are a mixture of (by assumption) a few errors, confounding effects, and direct causal links. We next discuss situations in which the conditional independence observed relative to skeleton edges can be used as evidence for direction.

#### Theorem 3.2.

Let *G* be a genetic variable and (*X*, *Y*) be an edge in *G*_*T*_. If 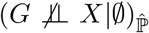, 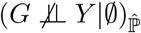, and 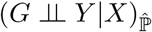, then *X* → *Y*.

*Proof.* Given the input and under the assumptions of no collider bias in the input dataset and local faithfulness, we can assume that there is a pathway *π*_1_ from *G* to *X* that has no colliders. We now look at the (*X, Y*) edge and consider four ways to orient it: (1) *X* ← *Y*, (2) 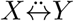, (3) *X ↔ Y* (i.e., a feedback loop), and (4) *X* ↔ *Y*. Options (1-3) imply that the new path from *G* to *Y* created by concatenating the new oriented edge to *π*_1_ will put *X* as a single collider. However, this pathway is unblocked given ∅ (by assumption) and should be blocked conditioned on *X*, a contradiction. We therefore have only option (4) remaining

Four important issues should be considered if we are to use such analysis in practice. First, note that we did not assume that *G* ∈ *Nei*_1_(*X*), and thus any variable associated with *X* can be used here. In fact, *G* may have direct associations with other phenotypes and the proof will remain the same. Second, notice that since we already observed 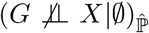 and 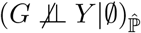 we can reasonably assume that there is enough detection power for associations in the examined local area of the causal diagram and thus use local faithfulness as evidence for blockage of a path. Third, when we analyze an edge (*X, Y*) in 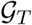 we exclude such variants that also satisfy the condition for the reverse analysis (*Y*, *X*). Such situations may occur when instruments are very weak, see simulated case 4 (**Supplementary Figure 5**). In addition, based on case 4, when we find such variants that satisfy the condition for both (*X, Y*) and (*Y*, *X*) this may indicate confounding by unobserved variables. Finally, in practice we check if 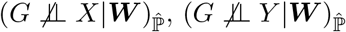, and 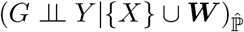, as by assumption ***W*** are exogenous (the proof remains similar).

### 3.7 DepEmerge: Emerging correlations

Here we make use of local faithfulness and observed collider bias in order to gain partial information about a pair. We consider “emerging” correlations: cases in which an a genetic variable *G* relative to a pair (*X*, *Y*), satisfies: (1) 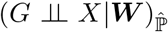, (2) 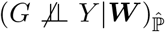, and (3) 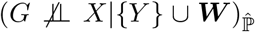. In words, the GWAS results indicate that the genetic variable *G* is independent of *X*, but becomes dependent once we add *Y* to the variables conditioned upon.

If the case above is detected we now have evidence for the existence of a path from *G* to *X* in which *Y* is a collider or a descendant of a collider and *X* is not. This covers two different cases: (1) *X* and *Y* are both results of a third variable *Z* that may be unobserved (confounding), or (2) *X* is a cause of *Y*. Since this information is ambiguous we mark these detected edges as *X*∗ → *Y*. If we have evidence for both *X*∗ → *Y* and *Y*∗ → *X* then we take it as evidence that either 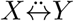 or there is a cycle that includes *X* and *Y*. The output of DepEmerge is interesting for two reasons: (1) it is another source of information to help us annotate skeleton edges, and (2) when many instruments support both *X*∗ → *Y* and *Y*∗ → *X* then DepEmerge reveals phenotype pairs for which we do not expect MR to work well even if there is prior knowledge about causal relationships between the phenotypes.

## 4 Practical considerations

### 4.1 Scoring each graphical analysis

The rules explained above are of two types. In the first we are looking for alternating significance of associations. This means that for each examined variant *G* and a pair (*X*, *Y*) we take two p-values *p*′ and *p*″ and ask, w.l.o.g, if *p*′ ≤ *p*_1_ ^ *p*″ ≥ *p*_2_. To summarize this analysis of a pair (*X, Y*) (e.g., summarizing all variants tested for this pair in section 3.4), we simply report the number and fraction of variants that satisfy the condition (out of all tested for the pair). The second analysis is much more similar to standard MR analyses. It is based on examining the association of a set of variants with a phenotype and performing a meta-analysis to summarize the evidence.

As a default analysis we run a standard meta-analysis based MR using the IVW method. In addition, for each analyzed pair, we examine the p-value distribution of the variants of the exposure with the outcome. Note that for inference of causal diagrams alone, instruments can point out potential flow of causal information without necessarily trying to estimate effects using a parametric model as MR methods do. In theory, under faithfulness, if *X* is a cause of *Y* then all instruments of *X* have and unblocked pathway with *Y* and should therefore be associated. Thus, we can simply ask if a set of instruments relative to *X*, *Y* is statistically associated with *Y* in a consistent way. We quantify this by computing an additional score: the proportion of non-null p-values, denoted as *π*̂_1_. This is a measure that is commonly used by FDR methods and it is estimated by comparing the observed distribution of p-values to a random uniform distribution [18]. Specifically, we use the local FDR method implemented in limma [19].

### 4.2 Interpretation of different signals

The analyses discussed above are fundamentally different. For non-skeleton edges we obtain a single unambiguous conclusion (once specific thresholds are set). The other analyses may provide different evidence for a pair *X*, *Y*. In practice, as we set *p*_1_ and *p*_2_ to be substantially different (e.g.,4-6 orders of magnitude) we assume that when detected, each evidence is reliable in pointing out existence of some path. Thus, when counting and examining patterns from multiple independent genetic instruments, a reliable comprehensive result can be attained. As an example in real data, we obtain evidence for both *Eosinophill* → *Asthma* and *Asthma* ∗→ *Eosinophill*. Thus, as a whole, the data has evidence for both direction and confounding, possibly by behavior or medication in this case. Nevertheless, we do note that the need for interpretation of such signals (possibly manually) and the fact that we do not directly estimate the FDR of the inferred networks are both limitations that can be addressed by future studies.

---

**Supplementary Figure 1.**
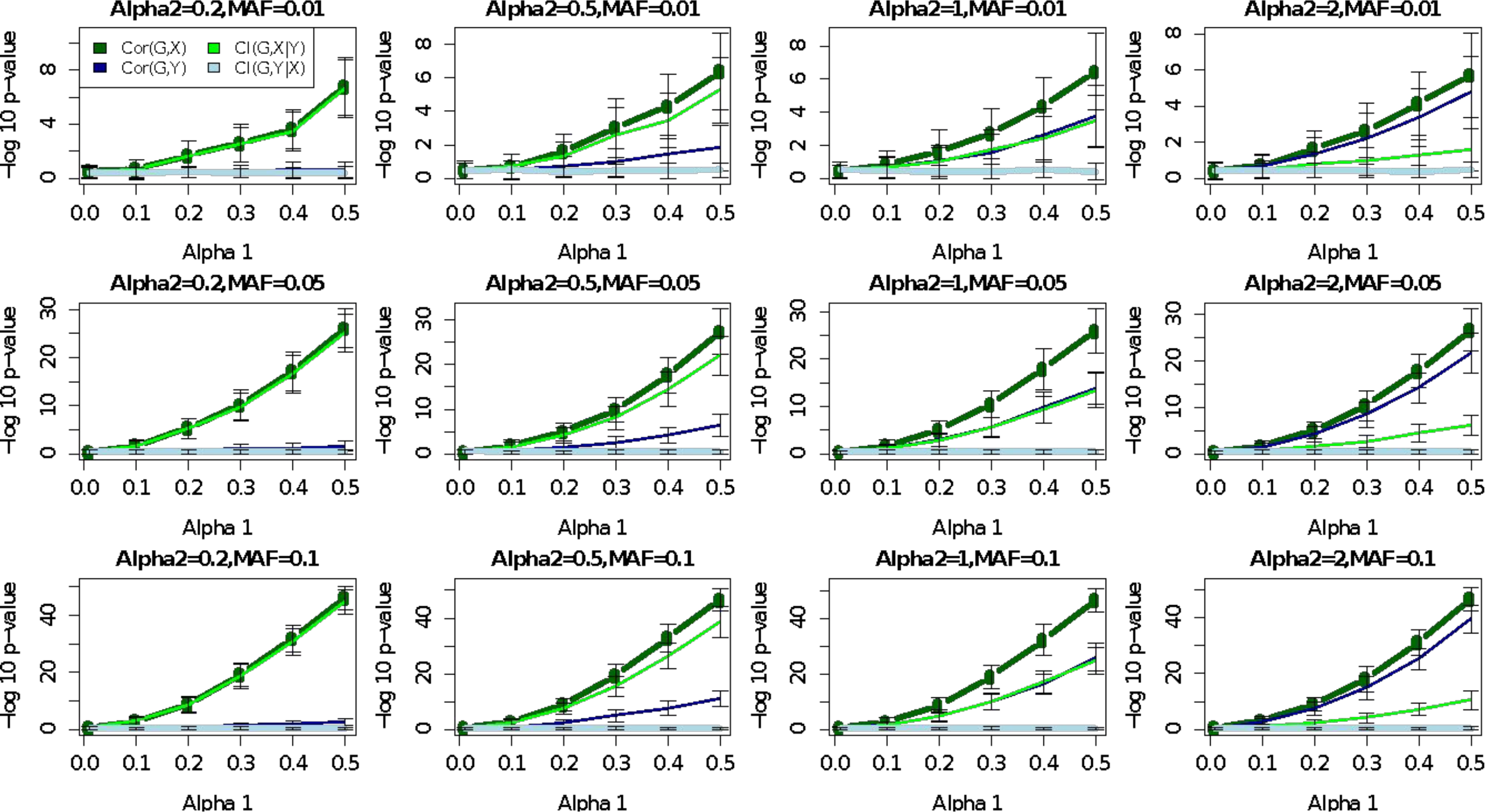
Simulation case 1. The graph structure is *G* → *X* → *Y*. *G* is a binary variable with different frequencies (“MAF”) whose linear effect on *X* is *α*_1_. *Y* is directly affected by *X* with *α*_2_. *X* and *Y* are continuous with standard normal noise. Each bar represents the results over 100 simulated datasets. Conditional independence was tested using linear regression or logistic regression.

**Supplementary Figure 2.**
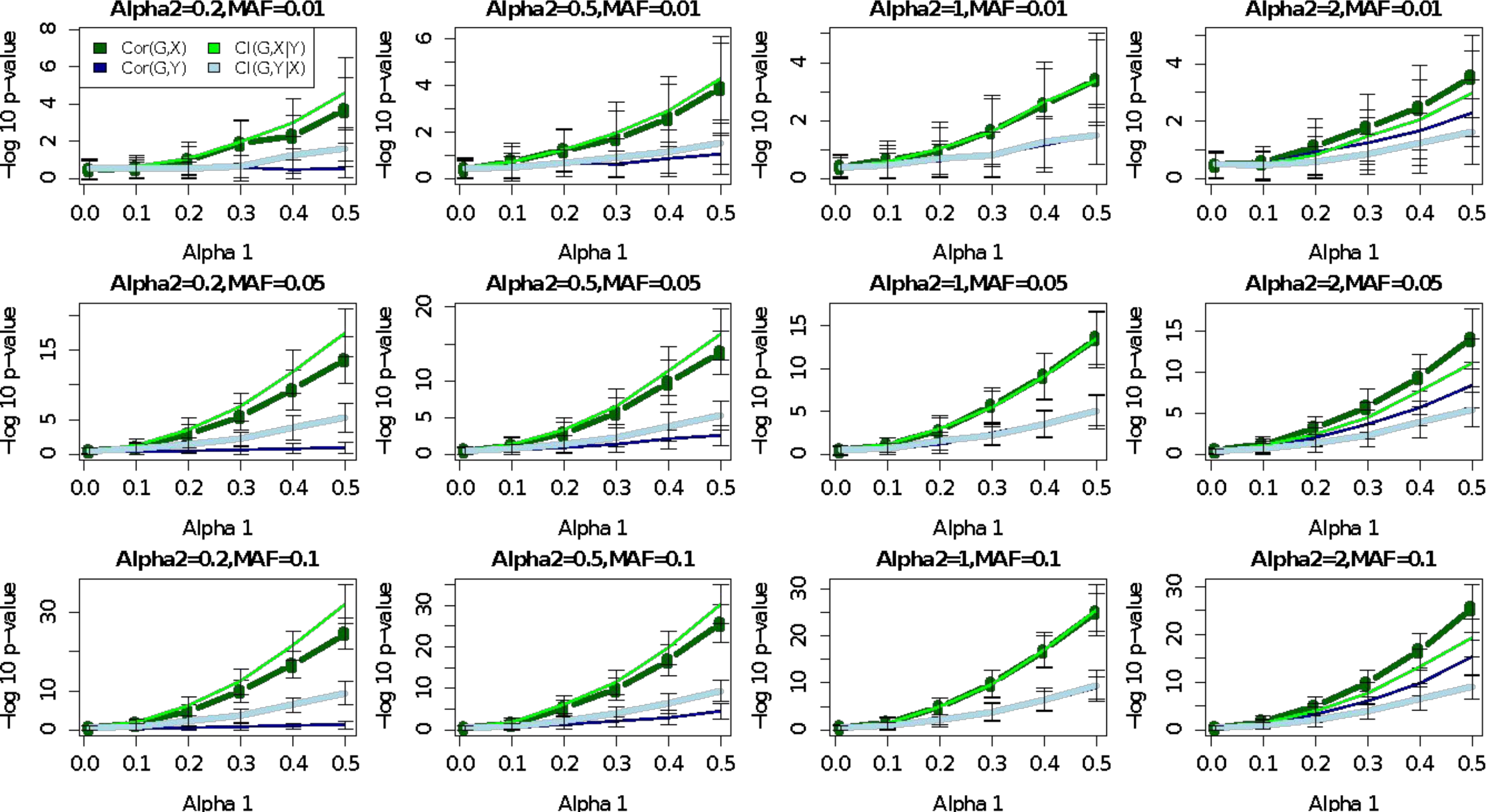
Simulation case 2 with strong confounding effects (*α*_3_ = 1). The graphical structure of the simulations is shown in **Figure 1A**. *G* is a binary variable with different frequencies (“MAF”) whose linear effect on *X* is *α*_1_. *Y* is directly affected by *X* with *α*_2_. *U* is an unobserved confounder with the same *α*_3_ effect on both *X* and *Y*. *U*, *X*, and *Y* are continuous with standard normal noise. Each bar represents the results over 100 simulated datasets. Conditional independence was tested using linear regression or logistic regression.

**Supplementary Figure 3.**
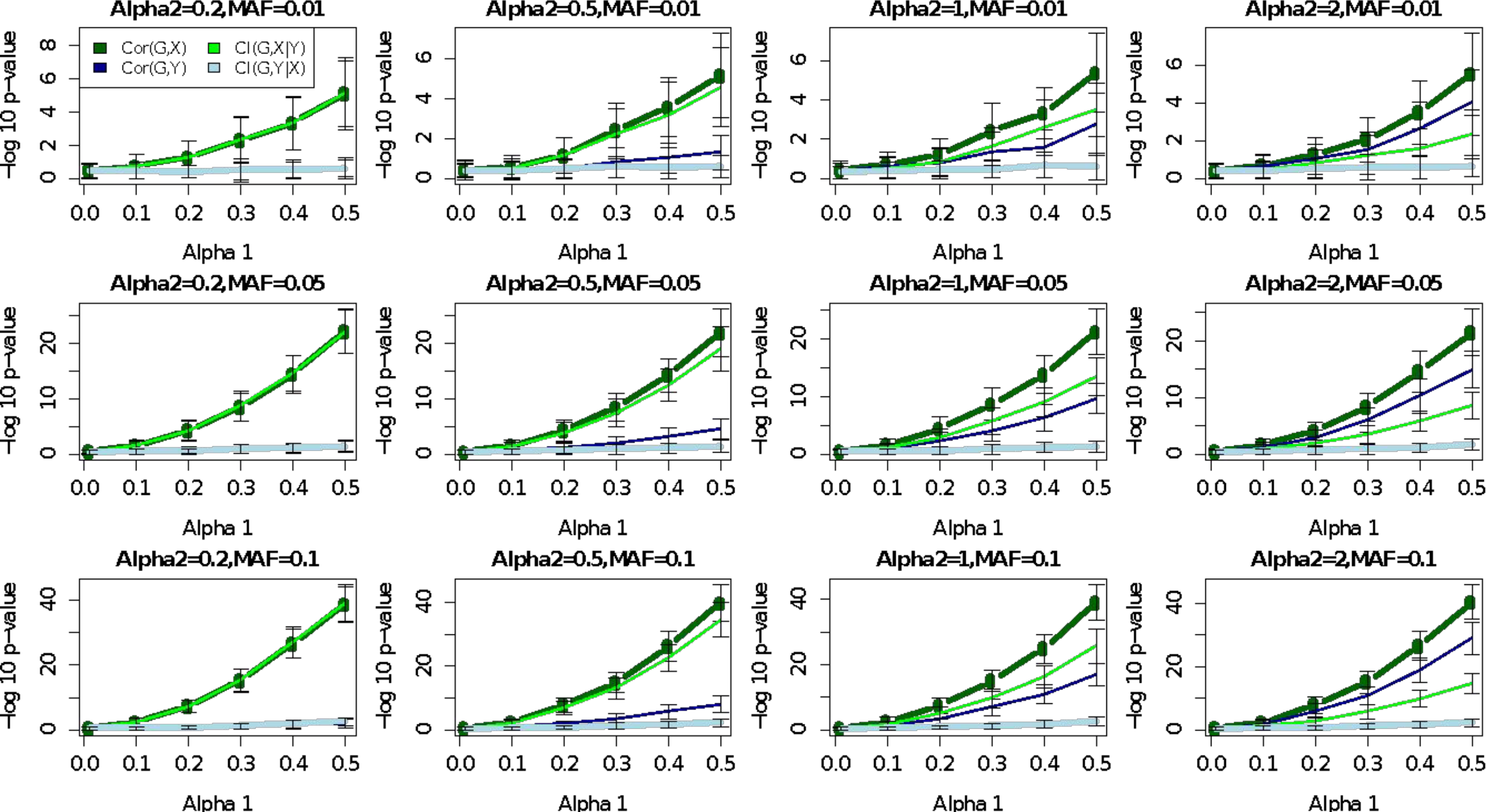
Simulation case 2 with weak confounding effects (*α*_3_ = 0.5). The graphical structure of the simulations is shown in **Figure 1A**. *G* is a binary variable with different frequencies (“MAF”) whose linear effect on *X* is *α*_1_. *Y* is directly affected by *X* with *α*_2_. *U* is an unobserved confounder with the same *α*_3_ effect on both *X* and *Y*. *U*, *X*, and *Y* are continuous with standard normal noise. Each bar represents the results over 100 simulated datasets. Conditional independence was tested using linear regression or logistic regression.

**Supplementary Figure 4.**
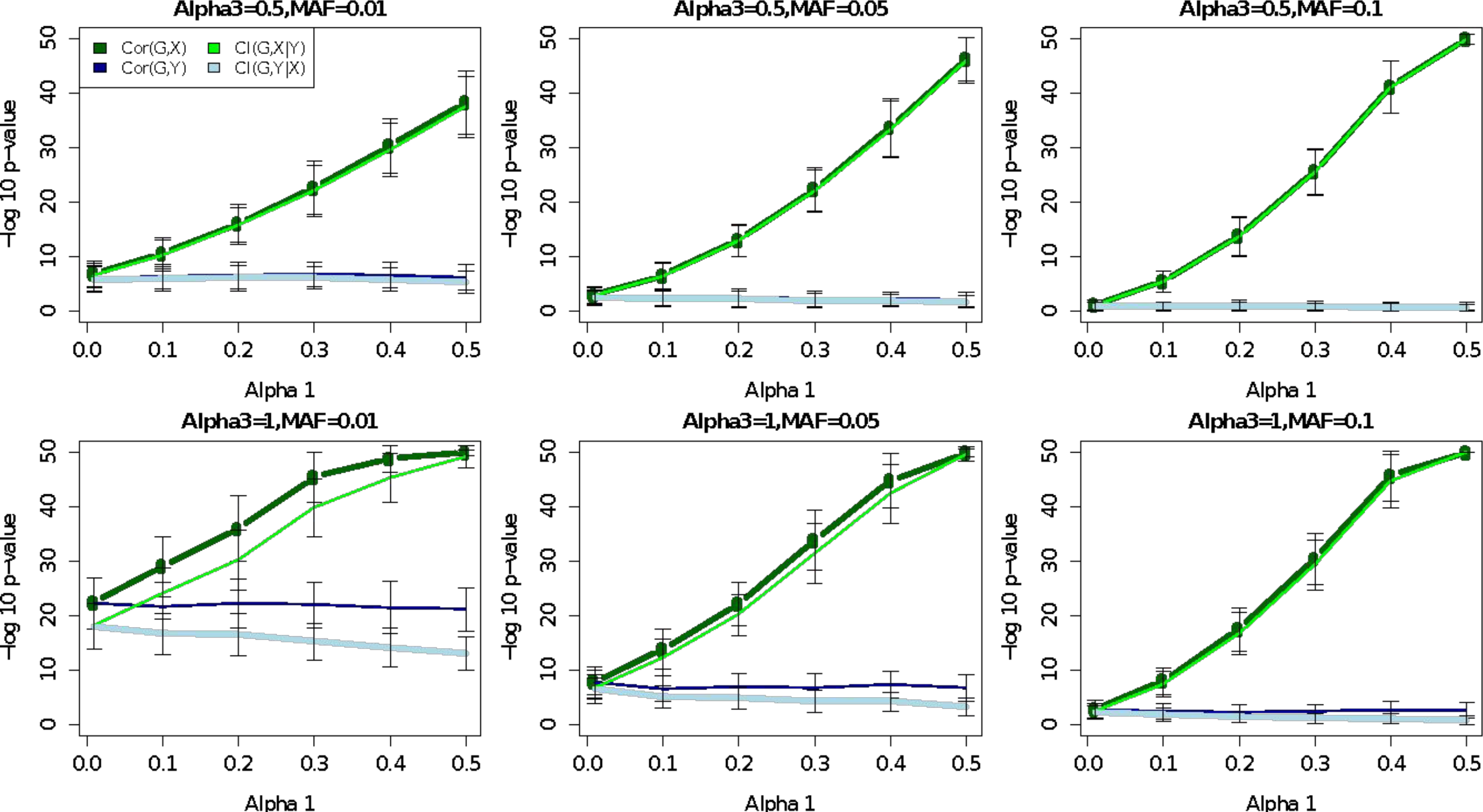
Simulation case 3, no direct effect with confounding (*α*_3_ = 0.5). The graphical structure of the simulations is shown in **Figure 1C**. *G* is a binary variable with different frequencies (“MAF”) whose linear effect on *X* is *α*_1_. *U* is an unobserved confounder with the same *α*_3_ effect on *X*, *G*, and *Y*. *U*, *X*, and *Y* are continuous with standard normal noise. Each bar represents the results over 100 simulated datasets. Conditional independence was tested using linear regression or logistic regression.

**Supplementary Figure 5.**
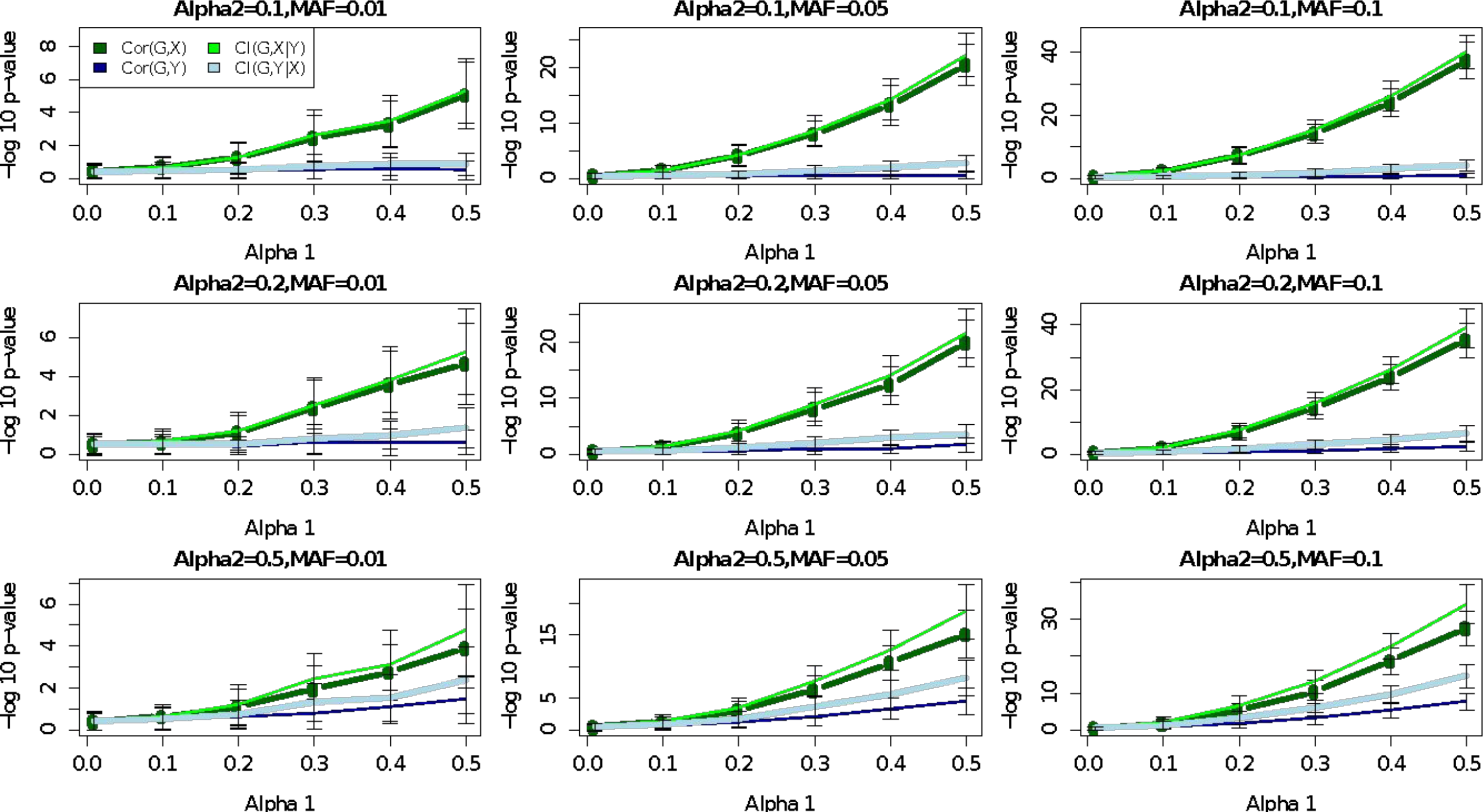
Simulation case 4, MR with a cycle (*α*_3_ = 0.5). The graphical structure of the simulations is shown in **Figure 1B**. *G* is a binary variable with different frequencies (“MAF”) whose linear effect on *X* is *α*_1_. *Y* is directly affected by *X* with *α*_2_. *X* is also affected by *Y* with the same coefficient *α*_2_. *U* is an unobserved confounder with the same *α*_3_ effect on both *X* and *Y*. *U*, *X*, and *Y* are continuous with standard normal noise. Each bar represents the results over 100 simulated datasets. Conditional independence was tested using linear regression or logistic regression.

**Supplementary Figure 6.**
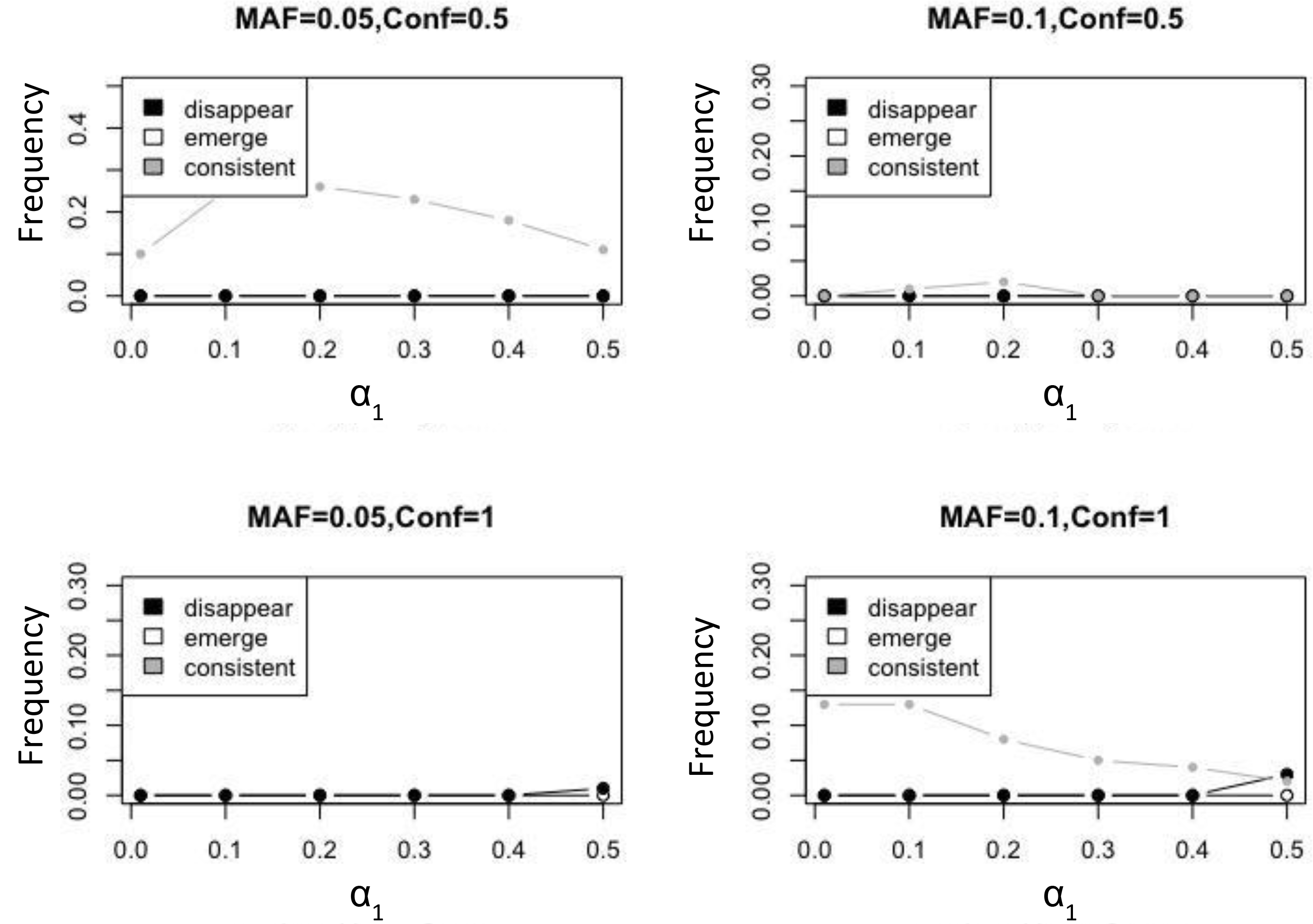
Emerging and disappearing associations are rare when all associations are confounded. The presented cases fit last cases of Figure 1. Each line presents the frequency of different patterns between the simulated genetic instrument and the phenotypes *Y* and *X*. In all counted cases *G* and *X* are significantly associated at *p* < 0.001. Disappearing: *G* and *Y* are associated but become independence when conditioned on *X* (*p* > 0.1). Emerging: *G* and *Y* are independent but become associated when conditioned on *X*. Consistent: *G* and *Y* are significantly associated with and without conditioning on *X*.

**Supplementary Figure 7.**
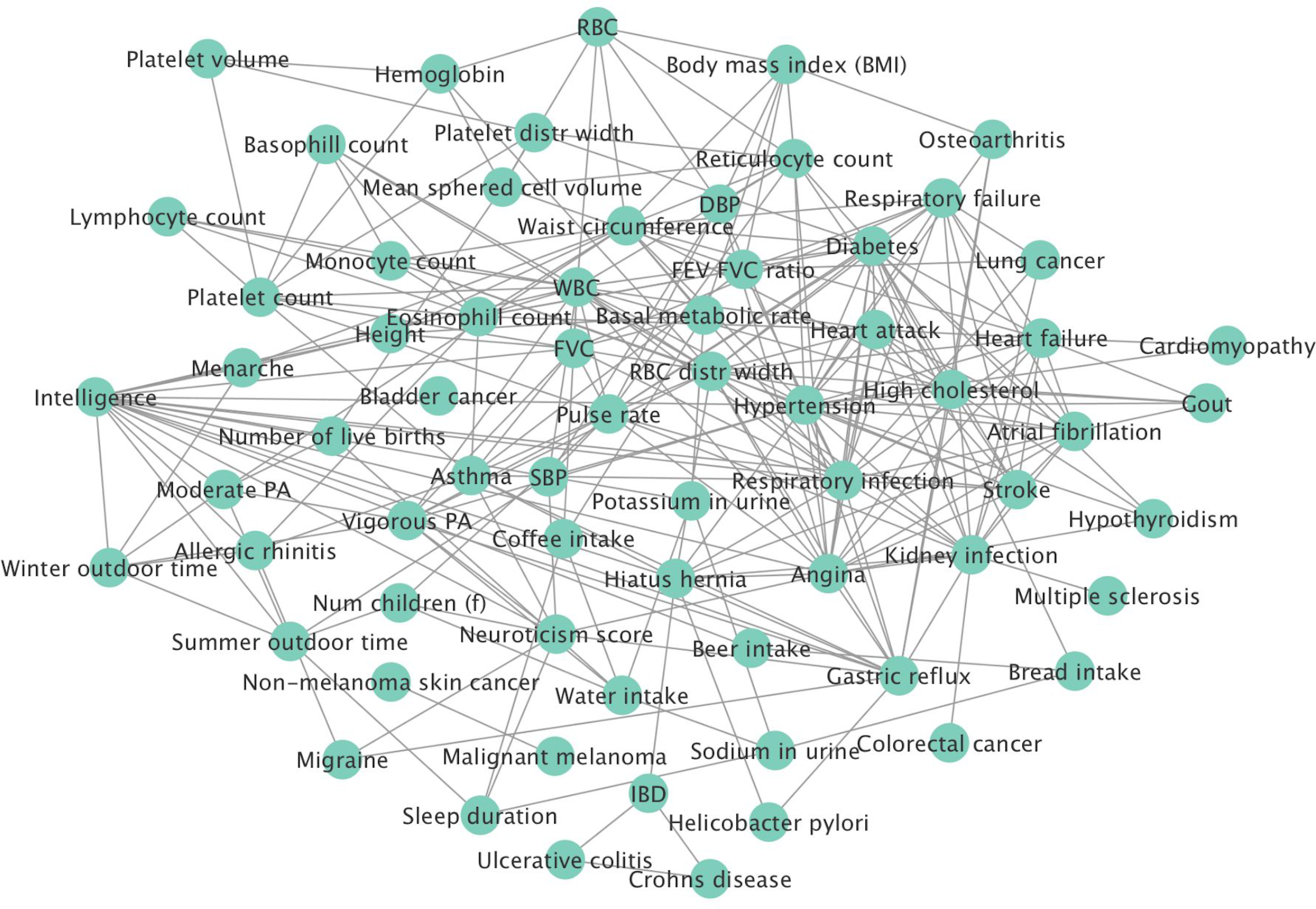
Inferred 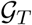 with *p*_1_ = 10^−6^. The edges represent phenotype pairs that remain associated at *p* < 10^−7^ when conditioned on other phenotypes. The graph structure can point out expected modules such as the dense area around heart attack.

**Supplementary Figure 8.**
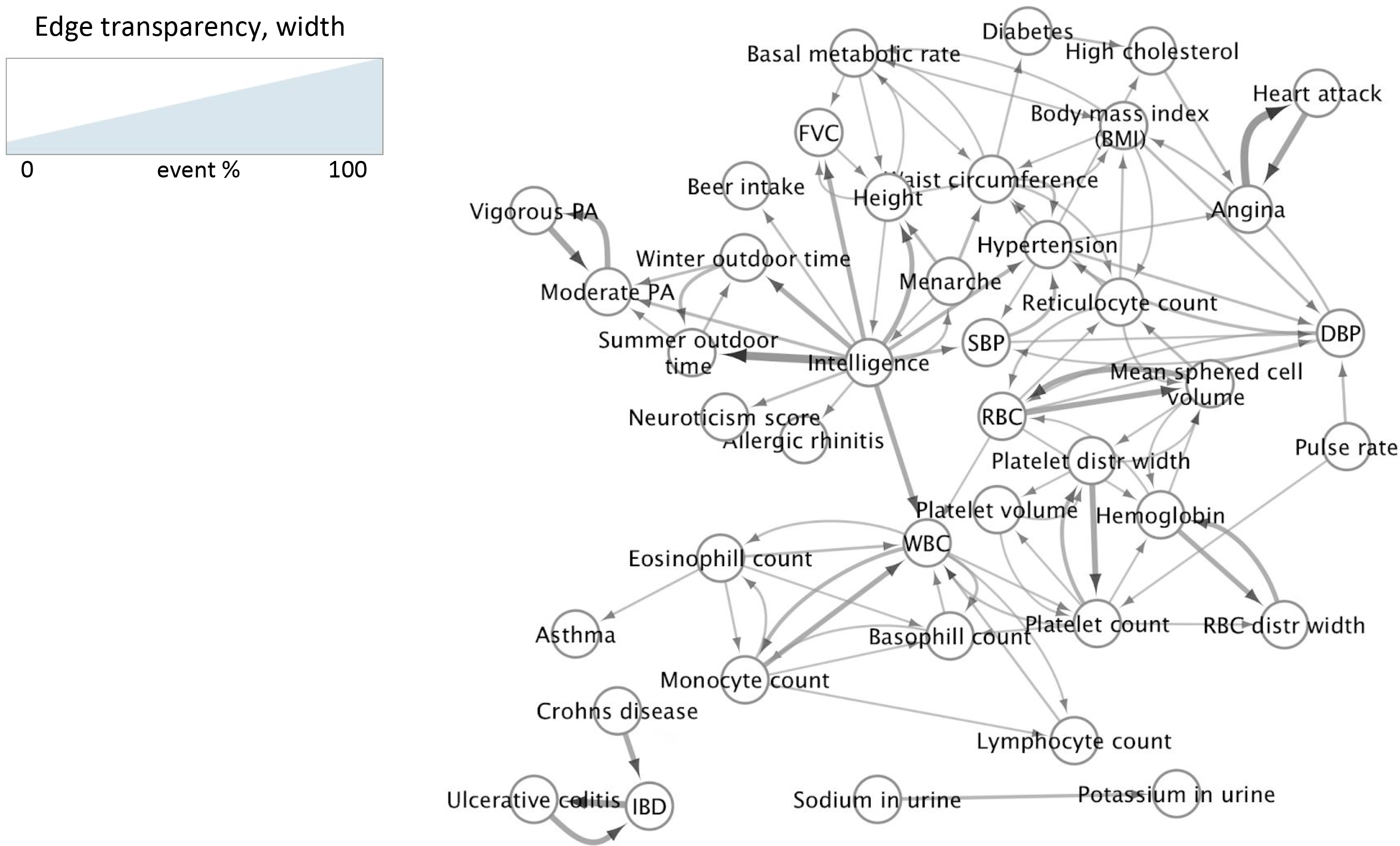
Inferred direct causal relations with *p*_1_ = 10^−6^, *p*_2_ = 0.001. The analysis reveals cycles, hubs, and clean directed paths. Intelligence, waist circumference, and whole blood count (WBC) are the main hubs.

**Supplementary Figure 9.**
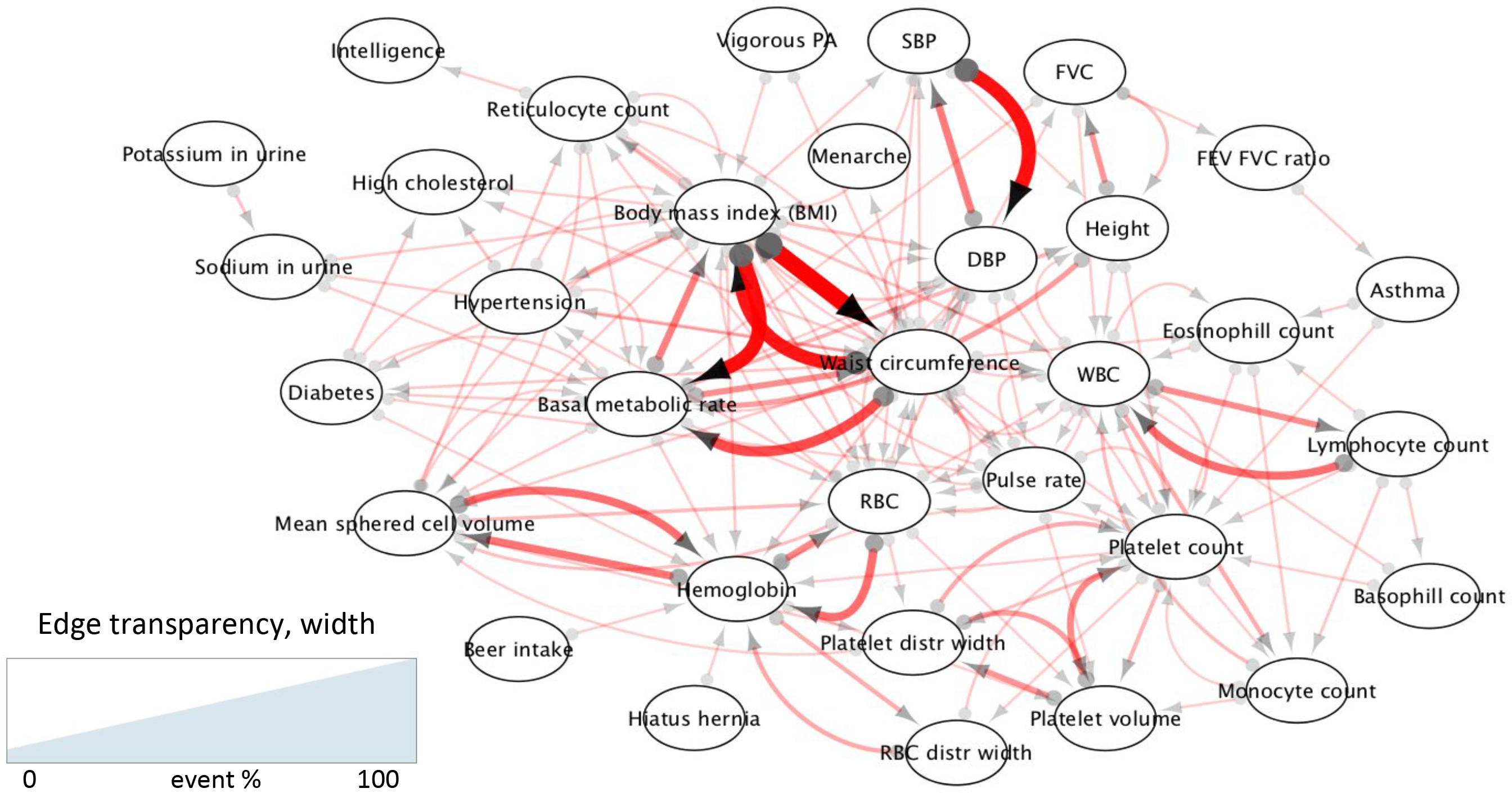
Inferred partial causal information with *p*_1_ = 10^−6^, *p*_2_ = 0.001. Each edge from X to Y represents evidence against a causal effect of Y on X suggested by the genetic variables. When there is an edge from X to Y and an edge from Y to X it serves as evidence for latent confounding. Most edges represent weak and partial information (edge consistency percentage scores are typically lower than 1%) but for some cases such as BMI vs. waist circumference the scores are high (> 50%).

